# A Compact, Standalone & Battery-Powered 3D Organoid-on-Chip System with Programmable Flow Control

**DOI:** 10.64898/2026.08.02.742326

**Authors:** Raviraj Thakur, Vaibhav Murthy, Samuel Olson, Emma Wolcott, Darya Budkina, Francis Anderson, Ting Zheng, Austin Wright, Jeremy Copperman, Luiz Bertassoni, Ellen Langer, Alexander E. Davies

## Abstract

Organoids-on-chip combine the 3D complex microenvironment and cellular composition of organoids with microfluidic flow, increasing nutrient-waste exchange and mimicking the contributions of *in vivo* interstitial and vascular flow. However, the widespread adoption of organoid-on-chip platforms is limited by the lack of incubator-friendly flow control systems. Existing approaches often rely on commercially available syringe or peristaltic pumps, but these are bulky, lack scalability, and present a significant barrier for clinical translation. To circumvent these issues, we present the Compact Active Perfusion Standalone Organoid-on-Chip (CAPS-OC) platform, a fully integrated and battery-powered microfluidic system capable of culturing organoids in active media flow. To achieve this, we introduce a novel low-power mechanism of pressure pulse generation using an off-the-shelf compact rotary actuator (CRA), and package it into a compact electromechanical assembly. This assembly provides timed pneumatic inputs to achieve programmable control of membrane-based peristaltic pumps, with ∼100 µL/hr dynamic range achieved on a custom microfluidic organoid chip. We biologically validated this system by culturing pancreatic cancer organoids derived from a Kras^LSL-G12D/+^; Trp53^LSL-R172H/WT^; Pdx1-Cre (KPC) genetically engineered mouse model. We found that our chip enhances proliferation and helps sustain a population of larger (>150 µm) organoids compared to standard dome based static culture. Additionally, through immunostaining, we observe that KPC organoids cultured in the chip show more aggressive PDAC phenotype with reduced nuclear expression of GATA6, whereas organoids in static culture retain less aggressive classical-like subtype. Finally, testing of a RAS inhibitor drug, daraxonrasib, on the KPC organoids on chip showed size-based sensitivity elucidating the impact of active perfusion on the drug diffusion kinetics. Altogether, we establish CAPS-OC as a valuable tool for the bioengineering community and for clinically translating organoid model systems.

## 1. Introduction

Advanced in-vitro cell culture systems such as organoids- and organ-on-chips (both abbreviated as OOCs here) are part of New Approach Methodologies (NAMs) that are currently being developed to reduce the burden of animal models in drug discovery and personalized medicine^1–3^. Often referred to as microphysiological systems (MPS)^4,5^, they have the unique ability to model highly spatially complex tumor or tissue microenvironments, using a combination of techniques from tissue engineering, cell biology and microfluidics. Fluid flow is an integral part of such device systems since dynamic culture often results in better cell growth, higher proliferation, and long-term maintenance owing to efficient nutrient delivery and continuous removal of waste products^6^. More importantly, static systems, both 2D as well as 3D, are diffusion limited and thus can fail to capture in-vivo physiology in terms of drug transport, clearance, and spatiotemporal gradients of signaling molecules. A recent study showcased how fluid flow enhances culture duration for breast cancer organoids over static 3D Matrigel domes^7^. It has also been shown by numerous studies that fluid flow is required to exert physiological levels of shear stress on cells and to activate specific pathways for accurate modelling of disease states.^8,9^

Despite these findings, static cell culture remains the primary technique likely due to standardized protocols and ease-of-use compared to flow-based systems. Off-the-shelf pumps such as syringe pumps^10,11^ or peristaltic pumps^12,13^ remain the popular choice to perfuse cell culture media through microfluidic chips, however experimental setup of these systems within cell culture incubators introduces substantial challenges. One such challenge is that not all tissue culture incubators have access ports for power cable and tubing feedthrough from the exterior to the pumps within, and such ports that are present create an opportunity for contamination of the cell culture. Simultaneously, commercially available syringe pumps are not designed for high humidity conditions and long term operation can damage them permanently. Additionally, external pumping requires a large volume of media due to large dead volumes making applications involving small volumes, such as drug screening, impractical. On the other hand, commercial OOC platforms are relatively expensive, often necessitating that academic research labs design and build their own systems. There is a clear unmet need for a robust fluidic technology that solves several of these challenges and enables researchers to incorporate fluid flow seamlessly.

Several alternatives to conventional pumping methods have been proposed for microphysiological systems which can be grouped into active and passive strategies^14^. Rocker based systems use mechanical tilting of chips to create hydrostatic pressure to drive fluid in a reciprocating manner^15–17^. This option is scalable, and different tilting rates and tilt angles can generate different flow rates. However, it lacks sophisticated control over flow rates, and the flow is bi-directional, which does not replicate physiology. Other active mechanisms such as electro-osmosis^18,19^ and electrokinetic methods have also been implemented in cell culture systems. While they require no moving parts, their application in cell culture is rather limited owing to difficulties of adoption in high conductivity media. Additionally, the applied electric field can disrupt cell growth, lead to uncontrolled phenotypic changes, and undesired changes to the pH of the cell culture media. Piezoelectric pumps^20,21^ have a thin membrane that can be deflected using an electric field to generate peristaltic flow over a wide variety of flow rates, but design, construction and integration of these pumps within microfluidic devices is challenging, and there in OOCs is limited. Moreover, piezoelectric pumps also require high operational voltage and thus sophisticated driver electronics which require deeper expertise in electrical engineering to design. Overall, these active methods enable precise flow control but still suffer from the need for peripheral hardware control systems.

Owing to engineering complexities of active methods, several passive pumping strategies have also been developed. Gravity based pumps use pressure generated by fluid columns to drive media through the devices^22,23^. Pumps based on capillary action or siphoning effect to drive media using porous materials such as cotton yarn resistor^24^, porous PDMS^25^, or paper^26^ have also been implemented in cell culture systems. However, none of these approaches are suitable for long-term experiments and suffer from a lack of on-demand programmable control. Additionally, these pumps cannot maintain uniform flow rate as driving pressure difference changes continuously. Alternatively, the use of bio-actuation of cardiomyocytes has been repurposed for pumping cell culture; however, development remains at a proof-of-concept level and won’t be applicable as a general-purpose mechanism due to cell culture media compatibility issues^25,27^. Collectively, these approaches highlight that existing microfluidic perfusion technologies either depend on bulky peripheral equipment, lack precise programmability at microscale flow rates, or are not readily compatible with long-term incubator operation, thereby creating a critical technological gap for compact standalone flow systems.

Here, we developed a *<u>C</u>ompact <u>A</u>ctive <u>P</u>erfusion <u>S</u>tandalone <u>O</u>rganoid-on-<u>C</u>hip* (CAPS-OC) platform integrated with a low-power, battery-operated pneumatic generator capable of fully standalone operation. The system combines a multi-chamber microfluidic chip containing six recirculating flow units—each consisting of an organoid well, media reservoir, and membrane-based peristaltic micropump. The pneumatic controller is designed with a novel electromechanical actuation mechanism in which a servo-driven compact rotary actuator generates programmable pneumatic pressure pulses to actuate on-chip pumps using a 4-phase peristaltic sequence. This architecture eliminates the need for external pressure sources while maintaining precise flow regulation and incubator compatibility. Although membrane-based on-chip peristaltic pumps have been widely demonstrated for lab-on-chip applications, their operation typically depends on external pneumatic and control infrastructure, including solenoid valves for timing, manifolds, pressure regulators, and centralized pressure and vacuum sources, which restrict portability and incubator compatibility^28,29^. To our knowledge, this is the first fully battery-powered platform that integrates pneumatic actuation, programmable media recirculation, and microfluidic organoid culture in a single compact system.

To demonstrate biological utility of the CAPS-OC platform, we cultured Kras^LSL-G12D/+^; Trp53^LSL-R172H/WT^; Pdx1-Cre (KPC) organoids, a well-validated organoid model system derived from genetically engineered mouse model ^30–32^ under varying conditions and perturbations. These organoids faithfully recapitulate the histopathological and molecular features of human pancreatic ductal adenocarcinoma (PDAC), including ductal morphology, characteristic oncogenic signaling, and the capacity for invasive growth, making them a clinically relevant model system for drug discovery^30,33^. The current standard for culturing KPC organoids is a static culture, where KPC cells are suspended in dome-shaped droplets of Matrigel on standard tissue culture plates, and allowed to self-assemble to form organoids in the 3D supporting matrix. Our microfluidic system showed superior performance compared to static domes by promoting enhanced growth and proliferation under dynamic culture conditions. We evaluated the drug response of KPC organoids against a RAS inhibitor, daraxonrasib, on-chip using a combination of confocal imaging and a viability assay. In this setting, we demonstrated tunability of drug dosing kinetics by simply modulating the duty cycle of the on-chip peristaltic pumps, a type of spatiotemporal control not feasible with any static system. Altogether, these data establish the CAPS-OC platform as versatile tool for the bioengineering community to integrate physiologically relevant culture conditions with tunable drug delivery kinetics, enabling more predictive in vitro models for studying disease progression and therapeutic response.

## 2. Methods

### 2.1. Pneumatic pressure controller design & assembly

#### Electromechanical assembly

The pneumatic pressure controller is an electromechanical assembly that consists of three main categories, 1) compact rotary actuator (CRA)–servo subassembly, 2) custom fixtures and fasteners, and 3) mounting plate-enclosure assembly. The CRA-servo assembly consists of a compact rotary actuator (McMaster-Carr 6508K103), CRA mounting bracket (McMaster-Carr 6508K42), flexible shaft coupling (McMaster-Carr 6208K431), servo coupler (Servo City 4001-0025-0250), DC servo motor (Parallax Inc. 900-00360) and a corresponding mounting bracket (McMaster-Carr 6508K42). Pneumatic tubing with 1/16” OD, 1/32” inner diameter (VWR MFLX06406-60) is connected to the CRAs via barbed adapters (Norgren ELB10-M3). 3D-printed parts are outlined in Supplemental Figure S2. All parts were 3D-printed (Raise 3D N2, Irvine, CA) in PLA using a 0.2 mm nozzle and 20% infill. The CRA (with mounting bracket) is attached to the CRA mount (Fig. S2a) via four M3 screws with lock washers and nuts. The servo motor is attached to the servo motor mount (Fig. S2b) via four M4 screws with lock washers and nuts. A triggering arm (Fig. S2c) is attached to the end of the CRA axle for interfacing with the microswitch for rotation registration. A 3D-printed enclosure (Fig. S2d) houses all the internal components, and a custom microscope adapter (Fig. S2e) is used to hold the chip on the microscope setup. Laser-cut parts are shown in Supplemental Figure S3. All parts are laser-cut (Boss Laser LS1416, Sanford, FL) from 1/8” thick acrylic at 50 W, 15 mm/sec. CRA and servo mounts are attached to the mid plate (Fig. S3c) via M4 screws, lock washers, and nuts. Microswitches are placed in small slits adjacent to the CRA mounts, and servo cables are passed through the rectangular cutouts in the mid plate. The assembled PCB is placed on the opposite side of the mid plate via four M3 screws with 6 mm standoffs, lock washers, and nuts. The mid plate, with attached CRA-servo subassemblies and PCB, is placed into the 3D-printed enclosure and secured with M3 screws. Brass M4 thermally set threaded inserts are inserted into the enclosure using a soldering iron tip, and the top plate (Fig. S3a) and bottom plate (Fig. S3b) are attached to them with M4 screws. Pneumatic tubing is passed through the small holes in the top plate. Supplemental Fig. S4 and S5 provide details for assembling the system.

#### Driver electronics

The driving electronics for the system are laid out on a custom printed circuit board (PCB) for ease of assembly (Fig. S1a). The control circuitry is based around an Arduino Nano microcontroller, which is responsible for digital and analog input and output, as well as state control (Fig. S1b). JST connectors are used to deliver power and communications from the Nano to the two servos, as well as to deliver the limit switch signal back to the Nano for servo registration (Fig. S1c). The servo control signals (S1_CT and S2_CT) are connected to digital output pins on the Nano, which use pulse width modulation (PWM) to control servo movement. The servo feedback signals (S1_FB and S2_FB) are read by digital input pins on the Nano, where they are converted to a PWM modulation for determining servo angular location. The registration signals from the limit switches (S1_RG and S2_RG) are connected to digital input pins on the Nano that are pulled up through software using onboard pull-up resistors. When the limit switch is closed, the input pin is grounded, allowing the Nano to detect that the servo has registered its position. A 3-way position switch and a push-button motion switch are also connected to the PCB through JST connectors (Fig. S1d). The position switch allows the user to select the maximum clockwise position (when SW_DN is shorted to ground), the maximum counter-clockwise position (when SW_UP is shorted to ground), or at the midpoint between maximum clockwise and maximum counter-clockwise position (when neither SW_DN nor SW_UP are shorted to ground, i.e., the switch is in the middle position). The motion control button controls whether the movement is paused or continuous. If the button is depressed (i.e., MN_CT is shorted to ground), the movement is continuous, and the 3-way position switch is ignored. If the button is not depressed, the movement is paused, and the servos move to the position indicated by the 3-way position switch.

The system can log temperature and humidity data over time, enabled by the inclusion of a DHT11 temperature/humidity sensor, a MicroSD writer, and a real time clock on the PCB (Fig. S1e). The DHT11 is supplied 5 V and ground, and its data pin is connected to an analog input on the microcontroller (DHT_DAT). The MicroSD card reader also receives 5 V and ground from the microcontroller and has four additional pins connected to digital I/O pins on the Nano: clock (CLK), data out (DO), data in (DI), and chip select (CS1). The real time clock is also powered by the microcontroller and additionally has its data (SDA) and clock (SCL) pins connected to digital I/O pins on the Nano. A power shunt circuit is included on the PCB to prevent the power bank from restarting due to a low current draw when the servos are stationary (Fig. S1f). This shunt consists of a transistor switch controlled by a digital output pin on the microcontroller which, when open, causes 5 V to be applied across two 150 Ω resistors in series. When the switch is activated, a current of ∼67 mA flows through the resistors and prevents the power bank from shutting off. The PCB also features an alarm LED powered by a digital output pin of the microcontroller which can be programmed to activate when a faulty state is encountered.

#### Firmware

The firmware consists of two library files (a C++ file and header file) that define classes and functions, and an Arduino file that employs these library files to operate the system. In the Arduino file, many parameters can be adjusted to suit individual experiments through variables at the top of the file. Examples include the speed of the servos (FWD_RPM, REV_RPM), the maximum angle that the servos are allowed to rotate through (THETA_ALT), and the duty cycle of the pump (DUTY_CYCLE_0, DUTY_CYCLE_1, DUTY_CYCLE_3). All of these factors impact the flow rate and maximum pressure generated by the system.

During an experiment, the firmware first registers the extreme travel angle of the CRA by rotating clockwise until the triggering arm activates the micro limit switch. Due to the high pressures of the registration step, it is necessary to leave pneumatic tubing disconnected from the microfluidic chip to avoid damaging the membrane fluid pumps. Following registration, the CRA will then move counterclockwise to the center of its travel (the zero-pressure point) and hold for 60 seconds, allowing time for the user to attach pneumatic tubing to the microfluidic chip. From the zero-pressure point, rotations clockwise will put positive (negative) pressure on the right (left) CRA output (as seen looking down the shaft of the servo toward the CRA). This allows for multiplexing two CRA inputs to four pressure outputs. For the peristaltic pump, a single output is used on CRA2 and both outputs are used on CRA1. The two outputs of CRA1 are attached to the first and last pump in the peristaltic arrangement (making them synchronous with opposite pressure polarities), and the single output of CRA2 is attached to the middle pump. Peristaltic flow is produced in four stages: 1) pull vacuum on the first pump to open it (CRA1 to clockwise target), 2) pull vacuum on the second pump to open it (CRA2 to counterclockwise target), 3) close the first pump and open the last pump (CRA1 to counterclockwise target), and finally 4) close the middle pump to push out the liquid (CRA2 to clockwise target). The flow rate can be adjusted by changing the duty cycle (i.e., the delay between stage 4 and stage 1). Three duty cycles can be defined in the firmware and switched mid-experiment via the 3-position switch on the front of the enclosure. At the end of each pump cycle, the firmware records environmental parameters (time, temperature, humidity, number of cycles) and saves this data to a microSD card for later analysis. Both the firmware files (C++ library and Arduino code) are attached in the supplementary section of the paper.

### 2.2 Microfluidic chip fabrication

#### SU-8 mold fabrication

The SU-8 molds were fabricated using standard photolithography techniques documented in the literature. Mechanical grade 100 mm diameter native oxide silicon wafers (University Wafer, Boston, MA) were cleaned with acetone and isopropyl alcohol, dried with filtered nitrogen, and baked at 150 °C for 5 minutes to remove surface moisture. Electronics-grade hexamethyldisilazane (Fisher Scientific, Waltham, MA) was applied to the surface of the wafers using a spin coater at 500 rpm for 30 seconds to promote adhesion of the SU-8 photoresist. A 5mL puddle of SU-8 3050 (Kayaku Advanced Materials, Westborough, MA) was poured onto the center of the wafers, followed by a two-step spin coating process: 100 rpm/sec to 500 rpm held for 10 seconds, 300 rpm/sec to 3,000 rpm held for 30 seconds for a nominal final thickness of 50 µm. The wafers were then soft-baked at 95 °C for 15 minutes, followed by edge bead removal using a small amount of SU-8 developer on a cleanroom swab. A mask aligner (SÜSS MicroTec MA6, Garching, DE) was used to expose the soft-baked wafers through the transparency masks with an exposure dose of 150 mJ/cm^2^. A post-exposure bake was then performed at 95 °C for 3 minutes, after which the wafers were cooled to room temperature and immersed in SU-8 developer for 10 minutes with agitation. Wafers were then cleaned with isopropyl alcohol and dried with nitrogen. Finally, a hard bake was performed by ramping the wafers on a hotplate at 5°C/min to a temperature of 150 °C, holding for 5 minutes, and then allowing the wafers to cool to room temperature on the hotplate to avoid cracking and delamination. The surfaces of the wafers were then passivated with (tridecafluoro-1,1,2,2-tetrahydrooctyl) trichlorosilane (Gelest, Morrisville, PA) in a vacuum chamber.

#### PDMS chip fabrication

Polydimethylsiloxane (PDMS) in the form of SYLGARD 184 (Dow, Midland, MI) elastomer was used to fabricate the chips. The PDMS was mixed in a 1:10 ratio of hardener to elastomer, degassed under vacuum, and poured onto the passivated SU-8 molds. The molds were cured at 65 °C for at least 12 hours. The two PDMS membrane layers were made by spin coating the degassed mixture on the 100mm silicon wafers using a spin coater. A thin layer of SU-8 featureless photoresist was deposited using the protocol described above. The mixture was spun to a final speed of 600 rpm at the ramp rate of 100rpm/s for 50 seconds of total duration. The membranes achieved an average thickness of 120 µm as measured by the profilometer (Bruker DekTakXT, Billerica, MA), see Table 1 in the Supplementary Material for measurements. The wafers were then baked at 65 C for 4 hrs. Each of the PDMS membranes was bonded to the corresponding fluid and gas layers using an oxygen plasma for 1 min at roughly 100mTorr (Plasma cleaner, Pie Scientific). The bonded layers were then incubated for 12 hours for a permanent reversible seal and elimination of any residual moisture from the chip. Both the fluidic and gas layers with bonded membranes were then treated with UV light for 40 mins for sterilization before cell seeding.

Supplementary videos (Video S1 and Video S2) show the operation of entire system and microfluidic chip.

### 2.3 Engineering tests

#### Pressure and current consumption measurement

The output pressure of the CRA and current consumption was measured using pressure sensors (Adafruit MPRLS Ported Pressure Sensor Breakout, Product ID: 3965, Adafruit Industries) and current sensors (INA219 High Side DC Current Sensor Breakout - 26V ±3.2A Max - STEMMA Q, Product ID: 904, Adafruit Industries). An I2C expander board (TCA9548A I2C Multiplexer, Adafruit Industries) along with an Arduino Uno3 (Adafruit Industries) was used to monitor real time data from up to three separate pressure sensors and two current sensors simultaneously. A custom python script was written to acquire sensor data via serial communication using *pyserial* library with a sampling frequency of 20Hz.

#### Flow rate measurements

Due to the absence of more sophisticated techniques such as particle image velocimetry, flow rate measurements were limited to a simple volumetric method. Since the organoid microfluidic chip has a recirculatory loop with a single media reservoir, it could be used with such measurements. A separate microfluidic chip was designed which had an inlet and outlet connected by a pump integrated channel. The inlet was filled with a food color dye mixed with DI water, and the outflow was collected under a variety of conditions as described in the results section. The volume was withdrawn from the collection reservoir and then weighed to back calculate the total volumetric flow rate.

#### Long-term incubator testing

The pneumatic controller boxes were tested for longevity first on the bench and then inside the tissue culture incubator. The temperature and humidity data were retrieved from the on-board microSD card. In the firmware, a temporary variable was incremented with successful execution of a pumping cycle, and the final value of the variable was used to determine the number of pump strokes executed. The pneumatic boxes were tested until the rechargeable battery was fully drained.

### 2.6 Cell culture, Media and Chip/Plate seeding

#### Plating of KPC organoids

The KPC (Kras-LSL-G12D/+; Trp53-LSL-R172H/WT; Pdx1-Cre) organoid line WFP9612 was obtained as a generous gift from Dr. William Freed-Pastor at the Dana-Farber Cancer Institute. KPC organoids were removed from Matrigel using Corning Recovery Solution and dissociated with TrypLE Express for 10 minutes at 37°C. Cells were resuspended in organoid splitting media comprising Advanced DMEM/F12 (12634010, Gibco) supplemented with 1% HEPES (pH 7.2-7.5, 15630106, Thermo-Fisher), 1% Penicillin-streptomycin (15140122, Thermo-Fisher) and 1% GlutaMAX (3050061, Thermo-Fisher), counted using a benchtop cell counter (Countess) and resuspended in Matrigel356231, Corning to get a final concentration of 5000/10,000 cells in 12.5 μL of Matrigel per well of a 96-well plate or chip. Matrigel-organoid domes were allowed to cure at 37°C for 15 min following which organoid culture media was added, containing Advanced DMEM/F12, supplemented with 50% L-WRN conditioned media (CRL-3276, ATCC), 10% RSPO-1 conditioned media (SCC111, Sigma-Aldrich), 0.4% GlutaMAX, 1mM N-acetylcysteine (A9165-25G, Sigma-Aldrich), 10mM Nicotinamide (N0636-100G, Sigma-Aldrich), 0.5µM A83-01 (9001799, Cayman Chemical), 0.05µg/mL mEGF (315-09, PeproTech), 10µM SB202190 (10010399, Cayman Chemical), 500nM PGE2 (14010, Cayman Chemical). Organoids seeded for experiments were additionally stained with Hoechst (1:50000).

#### Chip seeding and controller box prep

Fluid and gas layers were aligned and reversibly assembled after the seeding step. 12.5uL of Matrigel with suspended organoids were pipetted in the open organoid well in the fluid layer followed by an incubation at 37°C for 30 minutes. After the Matrigel was solidified, the fluidic layer was aligned with the gas layer under a brightfield microscope using a 4X objective to ensure proper alignment between the valve seat of the fluid layer and the valve chamber in the gas layer. Next, the pneumatic controller box was switched on, and the assembled chip was connected to the CRA pneumatic feed via 1/16 PTFE microbore tubing after the initialization step was completed.

### 2.7 Confocal Imaging

Multi-day imaging of organoids was performed using a Nikon Ti-2 inverted microscope fitted with an Oko-Lab environmental chamber maintained at 37°C and 5%CO2. Large field of view images were captured with multiple stage positions per well using a Nikon Plan Apo 0.45 NA 10X air objective. Finer detailed images were obtained using a Nikon Plan Apo 0.80 NA 20X air objective and Fusion-BT camera (Hamamatsu). Image capture was automated using NIS-Elements AR software.

### 2.8 Live-dead staining and viability assessment of organoids

To assess the viability, organoids were plated in microfluidic chips and assayed 5 days post seeding. Live cells were assessed by Calcein AM, ethidium homodimer (Live-dead, L3224, Thermo Fisher Scientific) and MitoSpy (mitochondrial membrane integrity, 424807, Biolegend) staining as described by the manufacturer. A subset of wells in both systems were treated with 1% Triton-X 100 (BP151-500, Fisher BioReagents) for 1 hour prior to staining and served as a positive control for dead cells. Organoid survival was determined by measuring the fluorescence intensity of Calcein AM, ethidium homodimer and MitoSpy using NIS elements software (Version 5.42.022, Nikon).

### 2.9 Immunofluorescence

KPC organoids were fixed in 4% PFA (J19943-K2, Thermo Scientific) for 20 (plate) or 40min (chips) and permeabilized with 0.1% Triton-X 100 (BP151-500, Fisher Scientific). Organoids and spheroids were blocked with 3% BSA in PBS supplemented with 1% Tween-20 (blocking solution, AAJ20605AP, Fisher Scientific) for 1 hour, following which antibodies against Ki-67, HIF1a, Vimentin, b-Catenin, GATA6 were added in blocking solution and allowed to incubate overnight (two overnights for chips). The following morning, cells were washed with PBS supplemented with 0.5% Tween-20 and secondary antibodies (donkey-anti-mouse-Alexa-488, Thermo Fisher Scientific and donkey-anti-rabbit-Alexa-555, A-31572, Thermo Fisher Scientific) were added at a concentration of 1:200 for 2 hours (plate) or two overnights (chip) in blocking solution. Cells were then washed with PBS+0.5% Tween-20 and counterstained with Hoechst (1:100) for 10 min. Cells were rinsed with PBS and imaged the Nikon Ti2 automated microscope previously described for live cell imaging. Each sample was imaged at 20x magnification in confocal mode (described above).

### 2.10 Computational Methods

Main steps of the computational pipeline were adapted from Murthy et al., 2025 with minor changes described below.

#### Region of interest extraction

To extract subvolumes for reporter spheroids, coarse masks of reporter cell locations were extracted using Gaussian smoothing and intensity-based thresholding. Similar coarse masks of DAPI nuclear stain were used in KPC organoids. Each identified region was expanded 6um axially and 25um laterally to ensure inclusion of surrounding regions. The resulting multi-channel subvolumes were extracted as TIFF stacks for downstream analysis.

#### Nuclear segmentation

To obtain single-cell nuclear masks, we utilized a deep learning-based approach using the Cellpose framework. Representative 2D training images were generated from two channels (KPC, brightfield and nuclear stain) or three-channel (brightfield, reporter, nuclear stain) stacks from ROIs. Slices exhibiting meaningful intensity variation were selected for manual annotation, yielding a training set containing thousands of annotated images capturing diverse imaging conditions and timepoints. These annotated datasets were used to train a custom segmentation model initialized from the pre-trained Cellpose cyto3 model. After training, the optimized model was applied to experimental image stacks, performing full 3D nuclear segmentation. Full parameters and cellpose models are available on our provided code repository. Imaging and segmentation data for each ROI was stored within a custom h5 data file for ease of downstream analysis.

#### Volume tracking

Organoid volume was quantified from 3D confocal image stacks using ilastik^34,35^-generated segmentation mask. For each image stack, the organoid masks were extracted from the foreground mask channel and treated as a 3D binary volume. Connected organoid objects were identified using 3D connected-component labeling, and the number of voxels belonging to each object was calculated.

#### 2D segmentation

Two-dimensional segmentation of organoids was accomplished using the Precision Organoid Segmentation Technique (POST) algorithm. Raw organoid images were cropped to remove background regions before running through the POST algorithm. The effective diameter of each organoid was calculated from the segmented area and converted from pixels to microns using the standardized scale bars at 4x and 10x magnifications.

#### Immunofluorescence quantification

Percent positive Ki67 and GATA6 nuclei were quantified as the sum of TRITC voxel intensities across all z-slices within the nuclear mask. To classify nuclei as positive or negative, a two-component Gaussian Mixture Model (GMM) was fitted to the log-transformed total TRITC sum intensity or sumTRITC/DAPI ratio, pooled separately across all chips and plated organoids to account for differences in laser intensity and acquisition settings. GMM threshold was defined as the intersection point of the two fitted components. Organoids with fewer than 5 nuclei were excluded due to possible under segmentation of ROIs (nROIs=3 excluded).

## 3. Results & Discussion

### Electro-pneumatic pressure wave generation using compact rotary actuator (CRA)

The CAPS-OC microfluidic system presented in this paper consists of a multiplexed peristaltic pump-integrated microfluidic chip and a custom-built battery-powered pressure controller that pneumatically powers the chip. The overview of the entire system is shown in Fig. 1a along with a graphical 3D representation of the chip in Fig. 1b. The chip produces a close-loop recirculatory flow over each organoid well as shown in Fig. 1b (inset) using the on-chip peristaltic pumps. The rationale for choosing these pumps is that they have a small footprint because they are comprised of three microvalves in series and can be placed anywhere in the fluidic path. But as stated earlier, the peristaltic flow is only generated when they are actuated using precisely timed pneumatic signals. This requirement has been traditionally met by using off-chip solenoid valve arrays and pressure controllers. Our goal with this design was to overcome this limitation by developing a compact and standalone integrated system that still has active programmable flow control, but without needing any external peripherals. Fig. 1e shows an exploded view of main components of our electro-pneumatic assembly, whereas Fig. 1f shows the exploded view of the 6-plex organoid chip along with the bracket. Together, this system can be easily placed inside standard tissue culture incubators as seen in Fig. 1d and solves several pain points of existing pumps that require several peripherals outside as seen in Fig. 1c.

**Figure 1:**
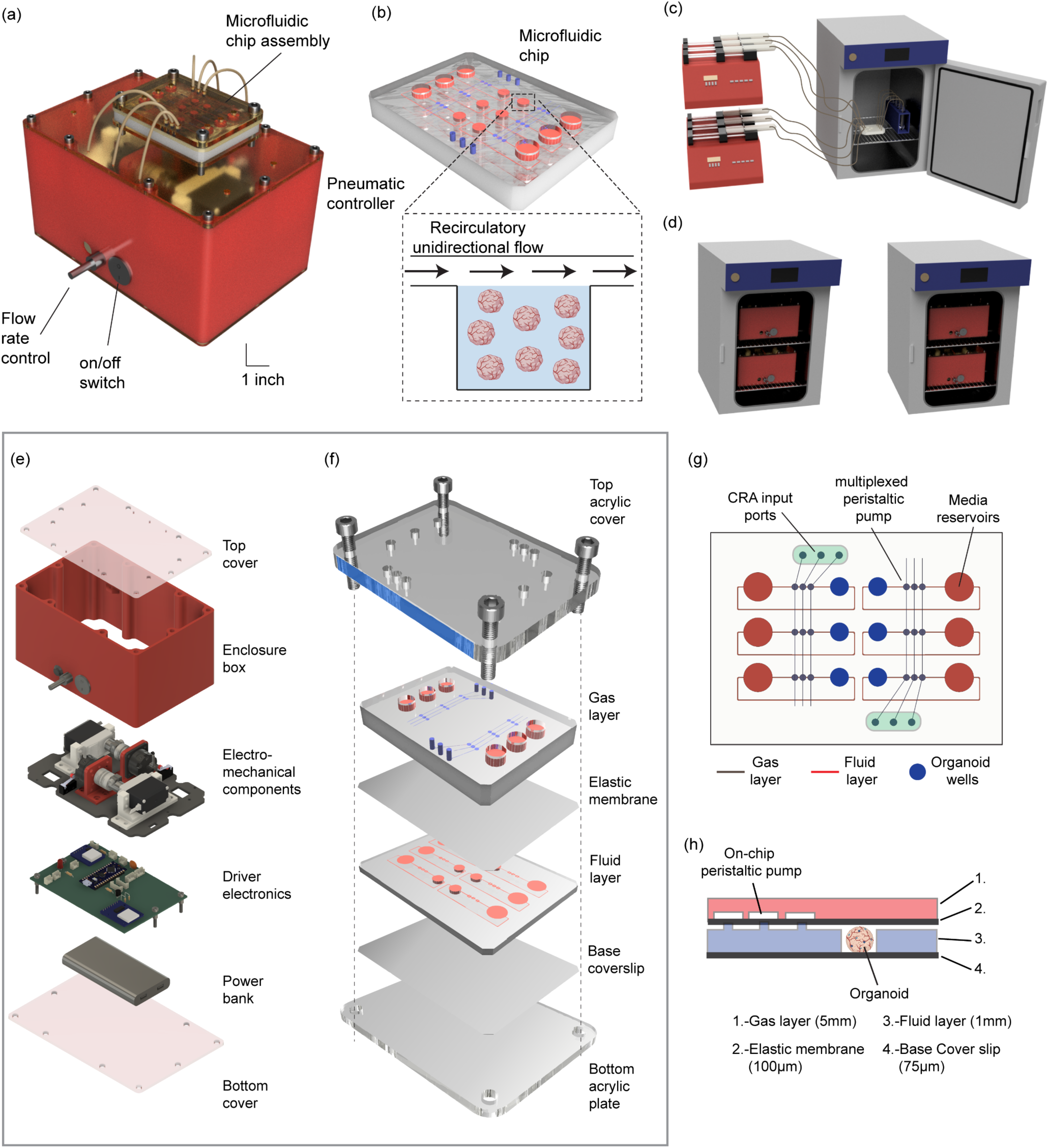
Overview of the proposed organoid on chip platform. (a) 3D overview of the integrated system consisting of the microfluidic chip and the battery powered electro-pneumatic controller. (b) Detailed 3D rendering of the microfluidic chip. (c) An illustration of a typical experimental setup for flow-based experiments using syringe pumps interfaced with microfluidic chips inside a tissue culture incubator. (d) Proposed compact and fully integrated flow-based platform highlighting elimination of external peripherals such as tubings and enabling scalability by stacking several units in the incubator. (e) An exploded view of the pneumatic controller assembly showing key components and parts. (f) An exploded view of the microfluidic PDMS chip showing multiple layers as well as the PMMA chip holder assembly. (g) A 2D schematic drawing showing spatial arrangement of the fluid and gas layers, specifically on-chip peristaltic pumps, media reservoirs and wells for seeding organoids. (h) A representative cross-sectional view of the pumps and organoid wells.

We hypothesized that CRAs can be repurposed for pressure generation when operated in reverse to their standard operational use, i.e. by mechanically turning their rotor to create a transient pressure pulse at their input ports. This pulse then can be fed to the micro-peristaltic pump on the chip to actuate the membrane valves and create peristaltic motion. The final form of the pneumatic pressure controller box is an enclosed assembly that has electromechanical subassemblies using CRAs, driver electronics, and a rechargeable battery resulting in a complete standalone system as shown in Fig. 1e. This system is designed with the purpose of supplying timely pneumatic inputs required for peristaltic flow as programmed in the firmware. The electromechanical assembly contains a 5 V direct current (DC) servo motor that rotates the central shaft of the CRA actuator. The motor was chosen such that it has sufficient torque to turn the CRA. The details of this subassembly are shown in Fig. 2a and 2b. A photo of the commercially available CRA is shown in Fig. 2c. An in-line flexible shaft coupling and a servo coupler were used to ensure maximum transmission of torque even for misalignments, as fixtures holding the servo motor and CRA are 3D printed with ∼100 µm resolution. A 3D printed arm was also attached on the other end of the CRA shaft to visually identify its angular position and to activate limit switches for rotation initialization.

**Figure 2:**
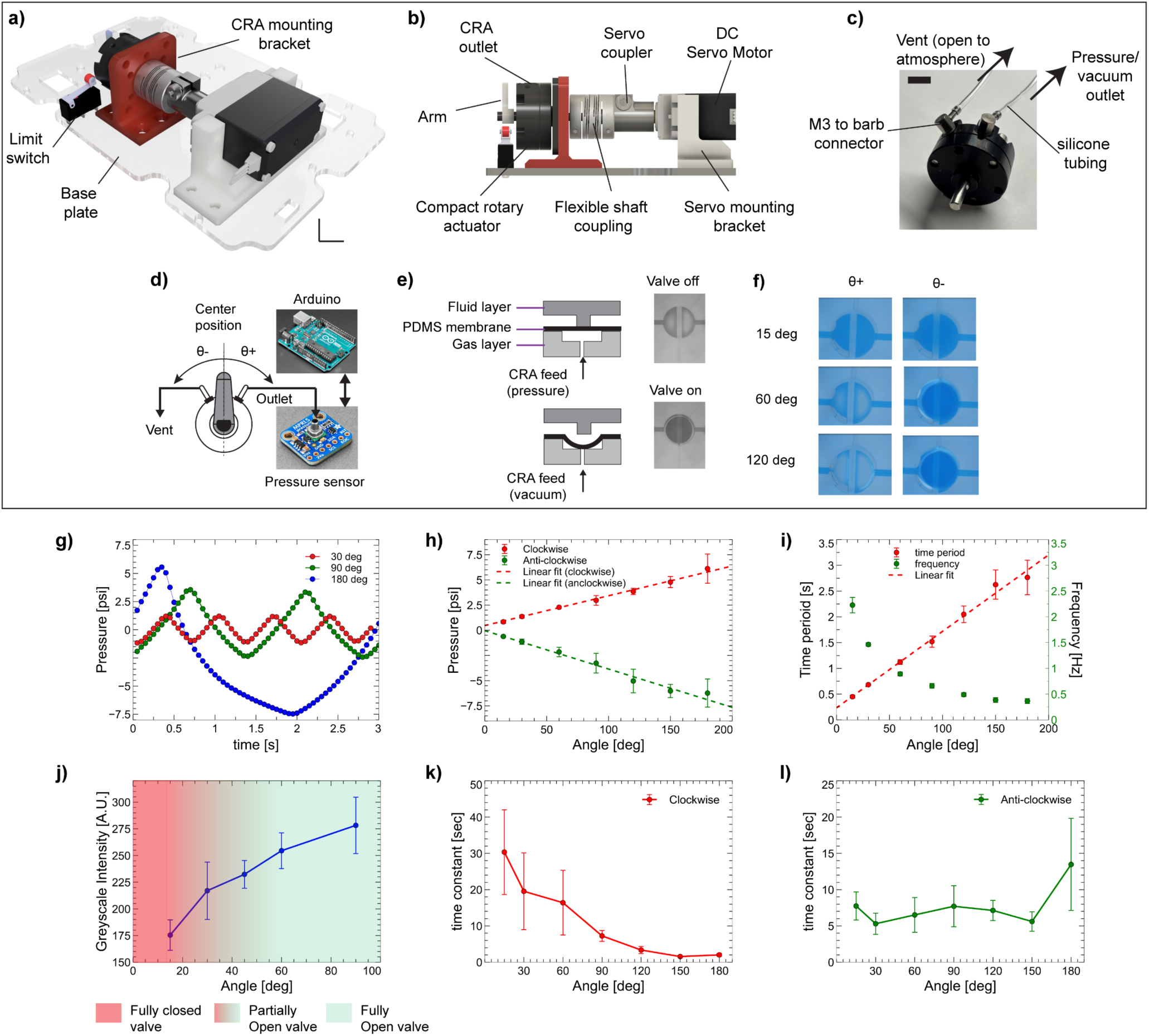
Compact rotary actuator (CRA)-based pneumatic actuation of a membrane microvalve. (a) An orthoscopic view showing the CRA-servo subassembly including mounting bracket, base plate and limit switches. (b) A detailed side view of the same assembly showing servo coupler, flexible shaft couplings and the DC motor. (c) Photograph of a commercial CRA with pneumatic output ports interfaced to silicone tubing via M3-to-barb connectors. (d) A schematic showing the CRA pressure output measurement using an arduino uno and pressure sensor in a dead-end configuration. (e) Side-view schematic and corresponding image of a normally closed microvalve of the three-valve on-chip peristaltic pumps. (f) Microscope images showing different angular displacements of CRA arm results in proportional microvalve opening. (g) Representative pressure output waveform as a function of different CRA rotation angles. (h) Peak pressure versus rotation angle, demonstrating an approximately linear response for both positive and negative angular displacements. (i) Output time period and frequency as a function of rotation angle. (j) Peak greyscale intensity measured as a proxy for valve opening. (k, l) Extracted time constants for pressure dissipation under clockwise and anticlockwise actuation, respectively.

To test our hypothesis, we characterized the output pressure as a function of the servo’s angle of rotation. For this measurement, one of the CRA ports was left open to atmosphere (vent), while the other was connected to a digital pressure sensor in a dead-end configuration, as shown in Fig. 2d. We observed that a clockwise rotation resulted in a positive pressure pulse at the other CRA port while an anticlockwise rotation resulted in a negative pressure pulse. Fig. 2g shows example pressure waves created by oscillating the servo between different extreme angles at constant speed. For the range of turn angles tested, we observed that the amplitude of pressure was linearly proportional to the rotation angle, for both clockwise and anticlockwise rotation as seen in Fig. 2h. The minimum achievable resolution of rotation was experimentally found to be 15°, as the servo could not reliably be moved in shorter increments. The lower limit of pressure was found to be around ∼1 psi for the 15° angle of rotation, and a maximum of ∼6.5 psi was achieved for a 180° angle of rotation. The rotational speed of the servo is held constant and shorter angles result in a higher frequency (Fig. 2g). These results confirm our initial assumption that an off-the-shelf CRA can be repurposed as a bidirectional pneumatic pressure source. Moreover, the linear relationship between rotation angle and output pressure provides a simple means of tuning the generated pressure through software alone.

Although the CRA was capable of generating measurable pressure pulses, the critical question for microfluidic operation is whether these pressures are sufficient to actuate membrane microvalves. Thus, we evaluated the valve state and state transitions as a function of CRA output. Fig. 2e shows the cross-sectional schematic of the microvalve along with the images of its two operational states, i.e. on or off. The output of the CRA was directly fed to the microvalve’s gas layer input port in the PDMS chip using silicone tubing. For leakproof connections, an M3 micro-barb connector was attached to the CRA. Fig. 2f shows example images of the microvalve for different angles of rotation. For angles less than 35 deg, the valve remained closed as evidenced by the closed valve seat, indicating there was insufficient vacuum to overcome the stiction force between the membrane and the valve seat. For angles of rotation greater than 60°, the valve was opened fully as confirmed by the intensity of the trapped dye shown in Fig. 2j.

Standard laboratory-grade pressure/vacuum sources can maintain a valve state by providing a continuous pressure or vacuum supply, whereas CRAs rely on a finite volume of mechanically compressed air and are therefore susceptible to pressure decay due to internal leakage over time. To characterize this behavior, we measured the transient pressure response of the CRA following a step change in servo position. As seen from (Fig. S6 left and right, supplemental section) the pressure builds as the servo turns and reaches its peak amplitude once the target rotation angle is reached and the servo stops. This peak is followed by an exponential decay in pressure toward 0 psi. We identified that this decay is due to the leakage in the CRA components and not from any of the interconnections. This is a somewhat undesirable characteristic; however, if the decay rate constant is sufficiently large, the valve state can still be maintained. Importantly, since peristaltic pumping relies on intermittent valve actuation, this leakage primarily limits maximum dwell time between two valve actuation signals, without affecting normal pump operation. Fig. 2k and Fig. 2l show the exponential decay time constants measured as a function of the step response at different angles, representing time required for a 63.2% drop from the peak. The decay rate for positive (clockwise) angles of rotation was vastly different than for negative (anticlockwise) angles of rotation. For positive angles of rotation greater than 120° the time constant was approximately 1 second, indicating a sharp decline from peak pressure, while positive angles of rotation below 60° resulted in time constants of ∼10 seconds. For negative angles of rotation, the time constant was found to be almost independent of the CRA turn angle and was consistently ∼5-10 seconds. These experiments conclude that in the current configuration, we can hold microvalve states for up to ∼16 seconds for angles less than 60° in the clockwise direction, and up to 5 seconds for all anticlockwise turns. Both of these time scales are substantially longer than the designed 300mS valve dwell time required during peristaltic pumping.

### 2/3 multiplexing CRA strategy for controlling 3-valve, 4 phase peristaltic pump

Having established that the CRA-servo assmebly can reliably generate pressures required for microvalve actuation, we next sought to minimize the number of CRA actuators required to operate the three-valve peristaltic pump. The on-chip pump in the microfluidic device consists of three sequential membrane microvalves that must be actuated in a specific sequence to generate peristaltic motion. A straightforward implementation would use three CRAs, each supplying one gas line to one valve. However, we leverage the inherent design of the CRA, which contains two chambers separated by a rotary vane, to simultaneously generate pressure and vacuum. As the total internal volume is fixed, rotation that compresses one chamber simultaneously expands the other, resulting in two output channels that are intrinsically out of phase. We exploit this characteristic to operate the three valves using only two CRAs instead of three. Reducing the number of actuators lowers system size and power consumption, while preserving the required valve actuation sequence. In this configuration, the two outputs of the first CRA are connected to the first and third valves, while the second CRA drives the middle valve, with its secondary outlet vented. Fig. 3a schematically illustrates how this arrangement enables a four-phase peristaltic actuation sequence. Inset images show the corresponding pump states visualized using food dye. Both CRAs begin from an idle position, denoted as phase [000], where 0 and 1 represent closed and open valve states, respectively. As the CRAs rotate according to a pre-programmed sequence, the four distinct phases generate peristaltic motion within the fluidic channel. Fig. 3b presents time-resolved measurements of output pressure, current consumption, and angular position during a single pumping cycle. Each valve state (or phase) is maintained for 300 ms, which is well below the system time constant described previously, ensuring reliable operation. The current consumption during state transitions is approximately 100 mA. In the angle–time plot, the red trace corresponds to the angular position of the servo driving CRA1, while the blue trace corresponds to CRA2. The total cycle time, measured via firmware, is approximately 2.5 s and is subsequently used to define the pump duty cycle for modulation of the net flow rate (see Table 2 in the Supplementary Material for duty cycle timing information). Following implementation of programmable control over the peristaltic pumps, we evaluated the tunability of flow rates using two primary control parameters: the rotation angle of the CRA/servo and the pump duty cycle. Flow rates were quantified using volumetric measurements as described in the Methods section. As shown in Fig. 3c, increasing the rotation angle from 15° to 60° resulted in a decrease in flow rate from ∼130 µL h⁻¹ to ∼40 µL h⁻¹, exhibiting an approximately linear trend. This behavior is consistent with reduced effective stroke volume at higher angular displacements, although further increases beyond 60° were avoided due to pressure dissipation effects noted previously. This limitation suggests a trade-off between mechanical actuation range and pressure transmission efficiency within the system.

**Figure 3:**
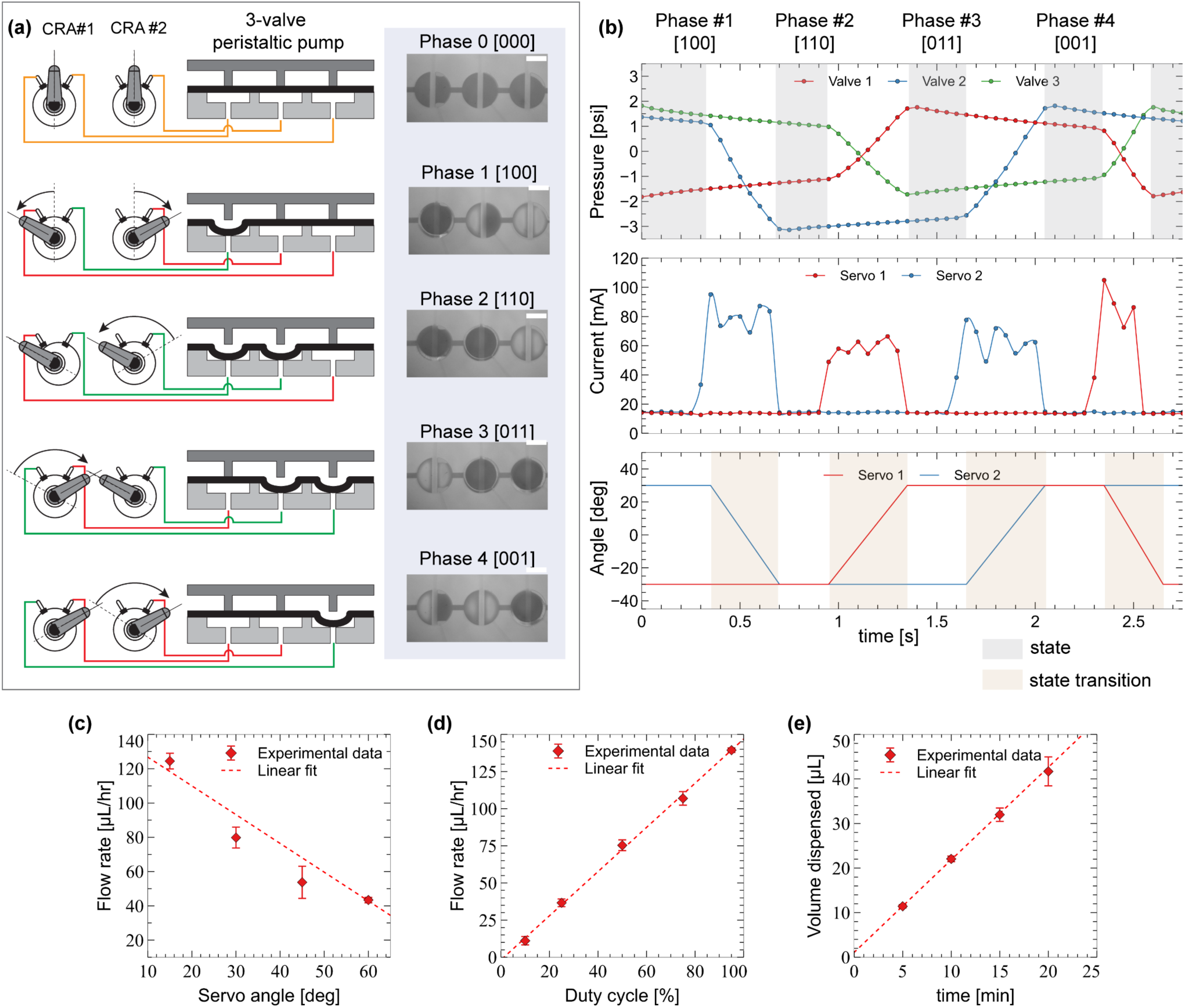
Multiplexed control and performance characterization of a three-valve on-chip peristaltic pump using two compact rotary actuators (CRAs). (a) Wiring diagram illustrating multiplexed pneumatic actuation of the three-valve peristaltic pump with two CRAs in a four-phase configuration. Inset: photographs of the pump at each phase during operation. (b) Time-resolved measurements of CRA pressure output, current consumption, and angular position. Each actuation phase is held for 300ms (shaded in light gray). (c) Cumulative volumetric flow rate as a function of CRA/servo rotation angle, showing an approximately linear relationship. (d) Volumetric flow rate measured at different peristaltic pump duty cycles, demonstrating a strong linear dependence. (e) Total dispensed volume over a 20 min interval as a function of time, showing linear accumulation consistent with stable pump performance.

In contrast, modulation via duty cycle provided a more effective and flexible means of controlling flow. Duty cycle, defined as the fraction of time the system remains in the active state within a given cycle, enabled a substantially wider dynamic range, spanning ∼10 µL h⁻¹ at 10% duty cycle to ∼145 µL h⁻¹ at 95% duty cycle as shown in Fig. 3d. The strong linear relationship observed (R² > 0.9945) indicates that flow rate scales predictably with actuation time, reflecting stable mechanical operation and minimal temporal drift. Importantly, this also demonstrates the controller’s ability to sustain repeated actuation cycles with varying dwell times without loss of performance, which is critical for implementing programmable flow profiles. From a practical standpoint, duty cycle control offers finer resolution and greater tunability compared to angular modulation, particularly for applications requiring low flow rates. Importantly, this mode of operation is fundamentally distinct from continuous low-flow perfusion, as it preserves higher instantaneous velocities and shear stresses during the pump on cycle, while reducing overall exposure through temporal modulation. This makes it well-suited for organoid culture systems, where precise regulation of nutrient delivery and waste removal is essential. Finally, the total volume dispensed increased linearly with time, as shown in Fig. 3e, confirming steady-state operation of the pump under constant fluidic resistance. This linearity indicates that the system maintains a consistent pressure head over extended periods, supporting its suitability for long-term perfusion experiments without the need for recalibration or external feedback control.

### CAPS-OC microfluidic design

There are two primary design considerations we used to develop a general-purpose flow-based microfluidic organoid chip. First, it should have multiple wells for seeding organoids to increase the overall throughput. Secondly, the chip design should allow imaging via confocal microscopy. The microfluidic chip design is shown in Fig. 1g. It consists of six flow cells, each consisting of a main reservoir for the cell culture media, an on chip microvalve peristaltic pump and a 4 mm organoid well all connected via a closed fluidic loop. The chip is comprised of 4 layers: a fluidic layer, a gas layer, and two thin PDMS thin membranes. Both membranes are bonded irreversibly to the fluid and gas layers. The membrane on the gas layer serves as the elastic layer for valve actuation while the membrane on the fluid layer serves as a base coverslip for imaging. The cross-sectional dimensions of the fluidic channel are 200 µm in width and 25 µm in depth. The organoid well is punched in the fluidic layer and sealed with a thin PDMS membrane at the bottom. The thin membrane allows use of high magnification confocal imaging. The chip has a total of six compartmentalized flow cells, permitting six different conditions of media or drugs per chip. The gas/control lines were designed in such a way that enables controlling all six different pumps with just one single CRA feed. The chip is housed in a PMMA bracket holder throughout the course of the experiments, shielding the chip from unwanted exposure to ambient surroundings and reducing the possibility of contamination. The inlet reservoirs are punched using a 10 mm biopsy and can hold up to 200 µL of media volume. Fig. 1f shows a 3D exploded view of different layers of this chip-holder subassembly, and Fig. 4a and 4b show an actual photograph of the microfluidic chip and the acrylic holder, while Fig. 4c shows the entire CAPS-OC system. Fig. 4d provides a block diagram of the entire system, showing mechanical, electrical and pneumatic connections between several key parts of this system.

**Figure 4:**
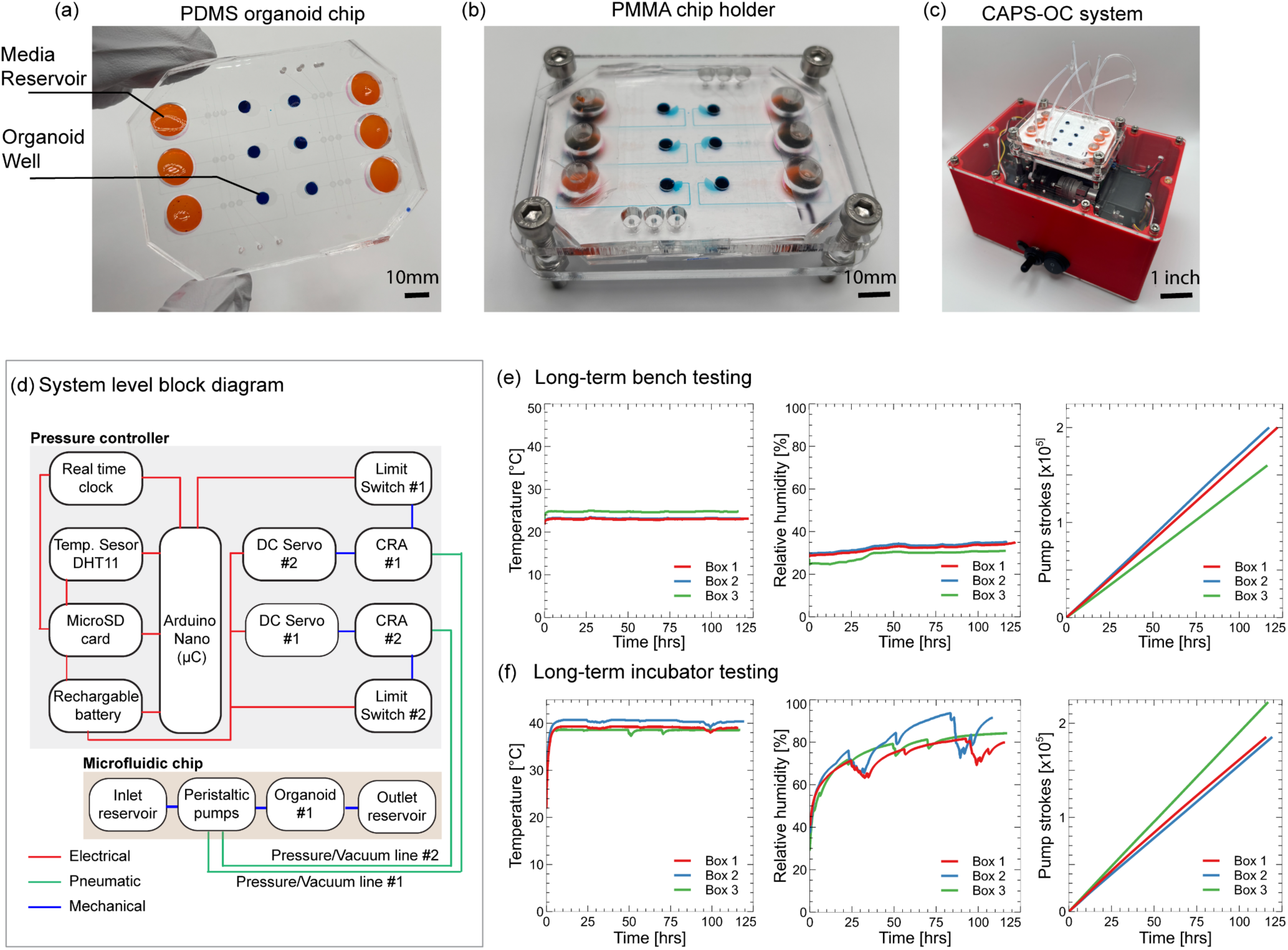
Bench and incubator performance of the microfluidic organoid-on-chip platform. (a) Photograph of an assembled PDMS microfluidic chip. Organoid wells are filled with dark blue dye, while reservoirs contain orange-colored water to visualize fluid distribution. (b) Microfluidic chip mounted in the holder assembly. (c) Photograph of the complete prototype, including the battery-powered electropneumatic actuator. (d) System-level block diagram illustrating the interconnection of electrical, pneumatic, and mechanical subsystems. (e, f) Long-term performance evaluation on the bench (e) and within a tissue culture incubator (f) over five days, demonstrating stable control of temperature, relative humidity, and peristaltic pumping.

### CAPS-OC maintains stable performance in tissue culture incubator

The pneumatic controller was evaluated under both benchtop and tissue culture incubator conditions to assess its thermal stability and to determine how reliably it can operate. Given the box contains active electromechanical components (DC servo motors), it was important to ensure that heat generation was within an acceptable range so as not to harm cells adversely. Additionally, the high humidity of the cell culture incubator can also corrode electronics and rotary linings over time. Hence, these tests were necessary before conducting biological validation of the CAPS-OC platform. Long-term testing showed consistent performance of the pneumatic controller in both environments, i.e. on the laboratory bench and in the incubator as outlined in Fig. 4e and f. The recorded temperature was stable around 23 °C on the bench, whereas it reached around 40 °C in the incubator. While this is somewhat higher than the incubator setpoint temperature of 37 °C, we believe this is internal to the box and does not affect cell culture performance in any way. Relative humidity levels reached up to ∼80% for the box in the incubator. Transient fluctuations are seen in both temperature and humidity traces correspond to brief disturbances caused by routine opening and closing of the incubator door. For these tests, a fully charged 50,000 mAh battery was used and was allowed to be drained fully to determine longevity of operation. Under a 95% duty cycle and a servo rotation angle of 60°, the system executed approximately 1.85 × 10⁵ pump strokes over ∼120 hours (5 days). For experiments requiring more than 5 days, the pneumatic box design allowed easy replacement of the battery by removing the bottom cover. These results validate the platform’s ability to sustain stable, long-term standalone operation under physiologically relevant incubator conditions.

### The CAPS-OC system enables better growth and higher proliferation compared to dome-based static cultures for KPC organoids

To demonstrate the biological utility of the platform, we assessed the effect of different flow rates in a PDAC organoid model. PDAC is a lethal human malignancy. It is characterized by a highly desmoplastic microenvironment ^36,37^ with dense matrix deposition, elevated interstitial fluid pressure^38^, and severely compromised vascular perfusion^39^, which collectively contribute to impaired drug delivery^39,40^ and aggressive tumor behavior ^41,42^. KPC organoids, derived from a well-validated Kras^LSL-G12D/+^; Trp53^LSL-R172H/WT^; Pdx1-Cre genetically-engineered mouse model ^30–32^ recapitulate the histopathological and molecular features of human PDAC, including ductal morphology, characteristic oncogenic signaling, and the capacity for invasive growth, making them an ideal system for modeling flow related biology in vitro ^30,33,43^. The current gold standard is a static culture where organoids are suspended in domes of ECM such as Matrigel. This method is fundamentally diffusion-limited and entirely lacks fluid transport, representing a significant gap between existing pre-clinical models and in-vivo disease phenotype ^44,45^. Recapitulating the dynamic fluid-mechanical context within the PDAC tumor in vitro requires a perfusion-capable platform with precise, programmable flow control.

The CAPS-OC platform uses PDMS as the primary chip material, and introduces some well-documented biocompatibility vulnerabilities that necessitate careful validation. For instance, PDMS has a well-established propensity to non-specifically absorb hydrophobic molecules ^46–49^ from cell culture media including growth factors ^47^, lipids^50^, steroid hormones^51^ and small molecules^52^ which could deplete critical paracrine signaling molecules when compared to a standard 96-well plate. Incompletely crosslinked PDMS can additionally leach into surrounding media and has been shown to have potential cytotoxic effects^53^. Additionally, while the confined geometry of the organoid well in the chip provides a design advantage, it could also affect organoid health. We tested these device-intrinsic effects by comparing the viability of KPC organoids cultured in a standard 96-well glass bottom plate and PDMS chips when subjected to high fluid flow (100μL/hr) 5 days after seeding. Viability was assessed using Calcein AM and Ethidium homodimer (EthDI) live-dead staining, while mitochondrial membrane integrity was measured using MitoSPY (Fig. 5a). We observed no difference in viability between the plate and chip conditions (Fig. 5b), confirming that the chip was biocompatible, and organoids were viable and healthy up to 5 days post-seeding.

**Figure 5.**
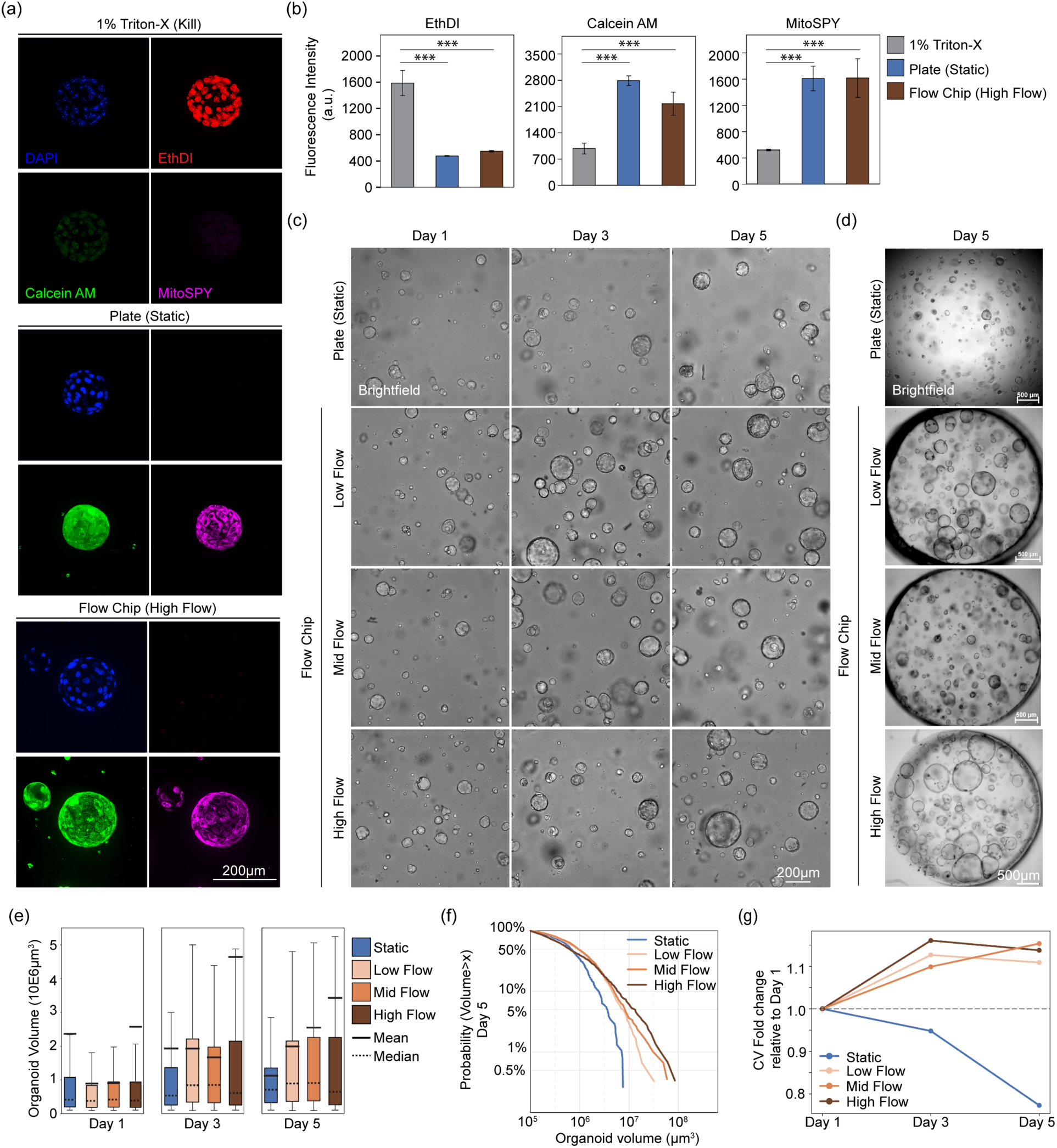
Biological validation of the microfluidic platform using KPC organoids. (a) Representative maximum intensity projection images of Ethidium homodimer (red), Calcein AM (green) and MitoSpy (mitochondrial activity, pink) 5 days post seeding of KPC organoids in static (middle) and high flow (bottom). Negative control organoids were treated with 1% Triton-X 100 for 1 hour at room temperature (top). Scale bar=200µm. (b) Fluorescence intensity quantification was performed from three independent experiments (N=3). Significance was determined by means of a Welch’s t-test (***-P≤0.001) on independent experiments. (c) Representative extended depth focused brightfield images of KPC organoids at day 1, day 3 and day 5 in static, low flow, mid flow and high flow obtained from 3D confocal microscopy. Scale bar=200µm. (d) Representative low magnification whole well extended focused images of KPC organoids at day 5 post seeding. Scale bar= 500µm. (e) Boxplot of volumetric measurements from 3D confocal imaging of KPC organoids in static (blue), low flow (peach), mid flow (orange) and high flow (brown) conditions. Solid black and dashed lines represent mean and median KPC organoid volumes respectively. Vertical lines denote inter-quartile range from greater than three independent experiments (N>3). n=2203 static, 3884 low flow, 4033 mid flow and 4483 high flow organoids across all timepoints. (f) Complementary cumulative distribution function (CCDF) of KPC organoid volumes under static (blue), low flow (peach), mid flow (orange) and high flow (brown) conditions at day 5. The y-axis represents the probability that a randomly selected organoid exceeds a given volume threshold (x-axis, log scale). n=349 static, 1215 low flow, 1960 mid flow and 1183 high flow organoids. (g) Coefficient of variation (CV) of organoid volumes over time under static (blue), low flow (peach), mid flow (orange) and high flow (brown) conditions. CV values are normalized to day 1 to illustrate the relative change in population heterogeneity across the experimental time course.

We next cultured KPC organoids in a range of defined flow rates within the CAPS-OC platform and assessed the extent to which controlled perfusion can drive flow-dependent responses in this clinically relevant cancer model. We quantified organoid volume across the static cultures and three flow conditions at three timepoints (day 1, day 3, day 5) using 3D confocal imaging (Fig. 5c and 5d). For all experiments, the volume of culture medium was kept constant between static and flow conditions since differences in volumes can potentially introduce several confounding factors such as disparity in nutrient availability, waste accumulation or concentration of secreted growth factors. The low-volume perfusion capability of CAPS-OC enabled this testing the effect of flow in much controlled manner, allowing direct comparison of the experimental conditions. Organoid volumes were calculated using brightfield masks by delineating organoid boundaries (against Matrigel) using a semi-supervised framework with the ilastik software package following manual training^54^ (Supplementary Fig. S7a). Volumes are therefore interpreted as segmented organoid volumes including the lumen, rather than cellular component walls or absolute geometric volumes. At day 1, all conditions displayed near identical volume distributions (Fig. 5c-e) confirming consistent and comparable baseline. Notably, flow induced a significant increase in volume by day 3 (Supplementary Fig. S7b, Fig. 5d and 5e) with median organoid volume increasing by 1.96-fold in combined flow conditions compared to 1.30-fold in static. By day 5, static organoids showed a marked narrowing of volume distribution, with an absence of a larger organoid subpopulation that was seen in flow conditions. In contrast, all three flow conditions maintained broad distributions with elevated medians compared to static. In particular, high flow conditions (100uL/mL) showed significantly elevated mean reflecting a small subpopulation of extremely large organoids (Fig. 5d and 5e). Statistical comparisons within groups and between groups have been summarized in Supplementary Table 6 and Supplementary Table 7. To characterize the behavior of the large-organoid subpopulation, complementary cumulative distribution functions (CCDF) were computed for each condition at day 5 (Fig. 5f). High flow maintained a shallow tail, retaining a measurable fraction of organoids at volumes approaching 10E8 cubic microns. Mid and low flow performed comparably with steeper declines beyond 10E7 cubic microns. Strikingly, static conditions produced no organoids exceeding 10E7 cubic microns by day 5, representing a lack of large organoids that were seen in the flow condition (Fig. 5f, Supplementary Fig. S7c). To determine whether flow altered population heterogeneity independently of size, the coefficient of variation (CV) was calculated on volumes for each condition across all timepoints. In all flow rates, the CV increased over the 5-day period suggesting that flow sustains population heterogeneity. In contrast, CV of static organoids decreased monotonically from day 1 to day 5 indicative of a converging population (Fig. 5g). Collectively, these data suggest that fluid flow is required to sustain volumetric heterogeneity of organoids with flow conditions producing large-organoid subpopulations that are not achievable through static cultures within the time frames we sampled.

The flow-rate dependent volume increase prompted us to ask if this expansion reflected an increase in cellular proliferation. To address this, we performed immunofluorescence staining for Ki-67, a canonical marker for cellular proliferation, in static and high-flow conditions on day 5 (Fig. 6a). Nuclear segmentation masks were generated using Cellpose3 applied to the DAPI channel of 3-D confocal z-stacks (Supplementary Fig. S8a). Ki-67 positive cells were determined using a two-component Gaussian Mixture Model (GMM) fitted to the total sum intensity in plate and high-flow conditions (Supplementary Fig. S8c). Organoids subject to high-flow conditions exhibited a significantly greater proportion of Ki-67 positive cells compared to static controls (Fig. 5b), suggesting that the volumetric differences are driven by increased proliferative activity rather than passive organoid lumen swelling or increase in cell size. For flow rate comparisons, GMM classification was performed on log-transformed TRITC/DAPI ratio, normalizing Ki-67 signal to nuclear DNA content (Supplementary Fig. S8d), with the low flow serving as the reference condition (Fig. 6f). Both Mid and high flow rates exhibited significantly higher Ki-67 positivity than low flow suggesting that a flow rate between 10-100μL/hr may optimally support organoid proliferation. These results are consistent with prior reports showing dynamic microfluidic environments promote organoid proliferation and tubularization in renal^55^ organoids and maturation of brain organoids^56^. In PDAC, elevated proliferation has been closely linked to loss of classical, progenitor-like tumor identity^57^ and is accompanied by epithelial-mesenchymal transition (EMT). To determine whether the flow-induced proliferative response was accompanied by broader phenotypic changes, we performed immunofluorescence staining for markers of EMT including Vimentin, and β-Catenin. We observed no qualitative differences between static and high-flow conditions (Supplementary Fig. S8b), implying that the flow-induced phenotypic response is not reflective of global de-differentiation.

**Figure 6.**
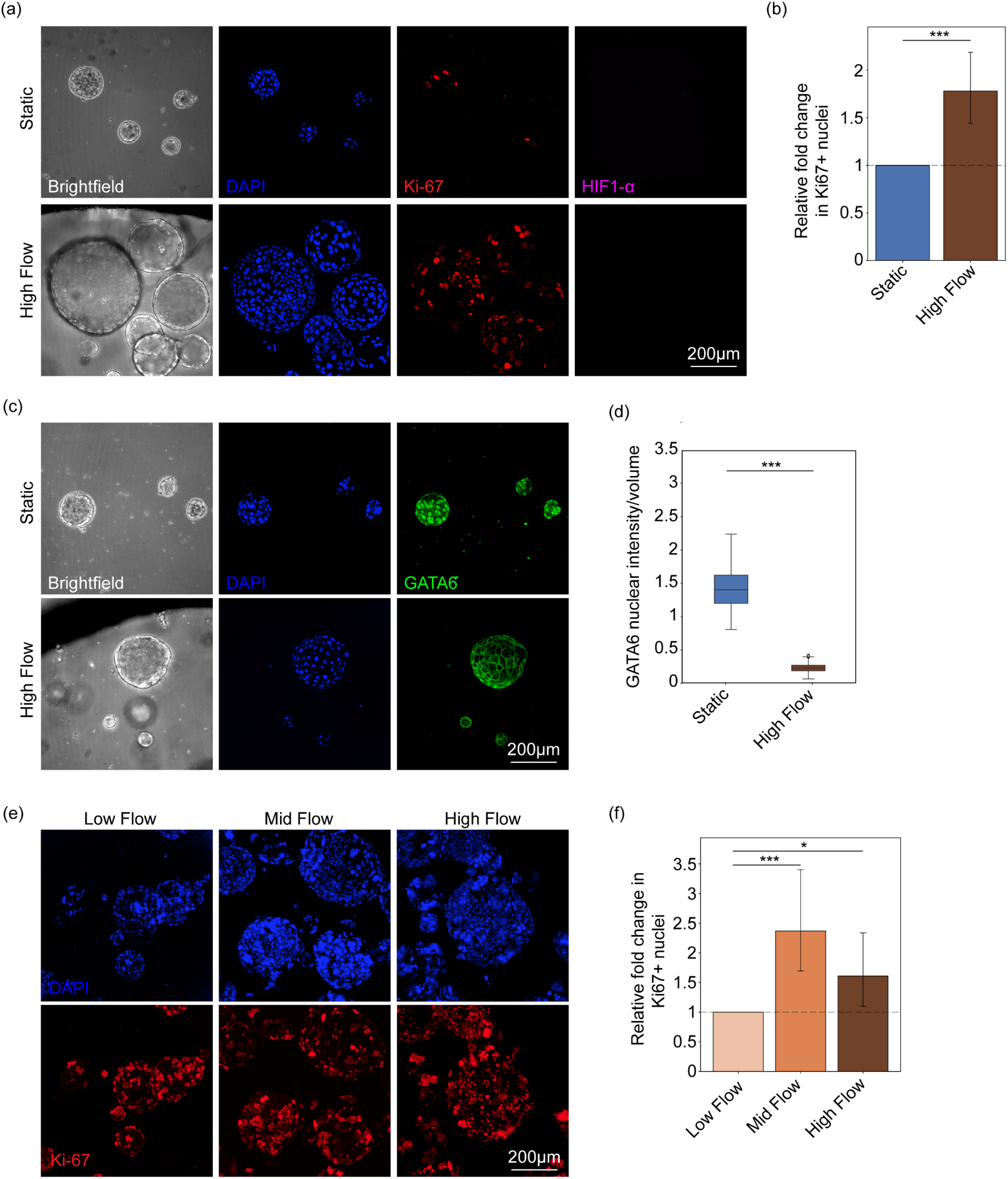
Immunofluorescent staining of KPC organoids on chip. (a) Representative maximum intensity projections of Brightfield (grey), DAPI (blue), Ki-67 (red), and HIF1-a (pink) staining in KPC organoids under static (top) and high flow (bottom) conditions imaged using 3-D confocal microscopy on day 5. Scale bar=200µm. (b) Boxplot showing quantification of relative percent positive Ki67 nuclei per organoid in static (blue), and high flow (100µ/hr, brown) conditions. Significance was determined using Wilcoxon rank-sum test (Mann-Whitney U) on per-ROI percent positive Ki-67 nuclei (***-P≤0.001) from two independent experiments (N=2). (c) Representative maximum intensity projections of Brightfield (grey), DAPI (blue) and GATA6 (green) staining in KPC organoids under static (top) and high flow (bottom) conditions imaged using 3-D confocal microscopy on day 5. Scale bar=200µm. (d) Boxplot of GATA6 intensity (GATA6 sum/volume) of nuclei per organoid in static (blue), and high flow (brown) conditions. Significance was determined using Welch’s t-test (***-P≤0.001) on at least two independent experiments (N=2). (e) Representative maximum intensity projections of DAPI (blue), and Ki-67 (red) staining in KPC organoids under low, mid and high flow conditions imaged using 3-D confocal microscopy on day 5. Scale bar=200µm. (f) Boxplot showing quantification of relative percent positive Ki67 nuclei in low flow (peach), mid flow (orange) and high flow (brown) conditions. Error bars represent 95% CI. Significance was determined using Wilcoxon rank-sum test (Mann-Whitney U) on per-ROI percent positive Ki-67 nuclei (*- P≤0.05, ***- P≤0.001)

Given that hypoxia is a well-established driver of aggressive, basal-like phenotype in PDAC and can itself promote EMT and increased proliferation, we sought to determine whether flow-induced changes were secondary to hypoxic stress. We stained for HIF1-α, and observed no significant differences between static and any of the flow conditions (Fig. 6a, 6e) suggesting that the observed changes may not be reflective of hypoxic stress. Acquisition of ‘basal’ phenotype in PDAC is marked by loss of nuclearGATA6, a central transcriptional regulator, where nuclear localization demarcates a differentiated ‘classical’ PDAC subtype^57,58^. The loss of nuclear GATA6 drives a transition to a ‘basal-like’ phenotype, which is associated with worse prognosis and resistance to therapy^58–60^. Notably, GATA6 expression and localization in PDAC organoids are not modulated by media composition alone, rendering it insensitive to routine culture perturbations^61^. Strikingly, immunofluorescence staining revealed a near complete loss of nuclear GATA6 in organoids cultured in high-flow conditions (Fig. 6c-6d), a pattern characteristic of the basal PDAC subtype. The absence of EMT markers in the context of GATA6 suggests that the flow-induced phenotypic response may represent an early or intermediate state of this subtype transition, preceding a full mesenchymal conversion. This is consistent with the notion that loss of nuclear GATA6 in PDAC can drive an epithelial-to-epithelial transition that precedes a full EMT program ^57^. The CAPS-OC platform was designed to decrease the effect of fluid-flow induced sheer stress, and the above proliferation and subtype transition towards a basal-like PDAC phenotype, suggests that these changes are driven at least in part by flow-related nutrient transport cues rather than hypoxic stress or global EMT.

### Microfluidic chip design enables user-friendly evaluation of targeted RAS inhibitor

To demonstrate the clinical and translational utility of the microfluidic platform, we evaluated the response of pancreatic ductal adenocarcinoma (PDAC) organoids to a targeted therapeutic. PDAC is predominantly driven by oncogenic RAS signaling, and since daraxonrasib is a direct RAS pathway inhibitor, we selected it as a representative agent to study drug response under physiologically relevant conditions. The KPC organoids were cultured and treated with daraxonrasib under three flow conditions (low, mid, and high flow) and compared against static dome cultures using conventional static 96-well plate. The organoids were treated with 20 µM of daraxonrasib on the 3^rd^ day post cell seeding. Each device contained paired control and treatment wells (n = 3 per group). For microfluidic experiments, organoids were maintained under high-flow conditions during the initial growth phase up to 3 days. On the dosing day, the duty cycles were changed to reflect low, mid and high flow conditions using the three-position switch on the controller box. Drug response was assessed using longitudinal brightfield imaging to quantify organoid size distributions every 24 h, complemented by an endpoint MTS viability assay after 72 h of treatment. Similar to previous experiments, the volume of media was kept constant between the static and flow conditions, so that total drug amount was same between the two groups.

For assessing viability using a standard MTS assay kit, our chip design permits adding MTS reagent directly in the input reservoir well as shown in Fig. 7(a). The on-chip pumps enabled continuous mixing of the MTS reagent with cells in small and controlled media volumes, and the effluent was easily accessible from the same media well after incubation. A custom 3D printed chip holder as seen in Fig. 7(b) was used to image the chips throughout the experiments under a confocal microscope. The compact and standalone characteristics of the platform are demonstrated in Fig. 7(c) where multiple CAPS-OCs systems are stacked in the cell culture incubator for KPC organoid growth and drug testing. Quantitative image segmentation analysis using kernel density estimation (Fig. 7d–k) revealed a progressive shift toward smaller organoid diameters in the treatment group across all conditions. No statistically significant differences were observed at baseline (T0), confirming experimental consistency for variables such as cell seeding density, Matrigel volume, etc. Over time, treatment groups exhibited both a reduction in mean organoid size and a narrowing of the size distribution, indicating suppression of growth and reduced heterogeneity. Notably, flow-dependent differences in drug response were observed on the terminal time point (72h). High-flow conditions produced the most uniform response, with a narrowest size distribution and absence of larger organoids (>150 µm), suggesting enhanced drug delivery and exposure. In contrast, low-flow conditions retained a subset of larger organoids after 72 h, indicating incomplete drug penetration or reduced transport. Mid-flow conditions exhibited an intermediate response, highlighting the role of perfusion in modulating therapeutic efficacy. Endpoint MTS viability measurements (Fig. 7f, h, j, l) showed consistent reductions in cell viability across all conditions, with approximately 25% viable cells remaining relative to the kill control (2% tritonX). Together, these results demonstrate that the organoid-on-chip platform enables flow-dependent evaluation of drug response and captures transport-mediated effects that are not observable in static culture systems.

**Figure 7:**
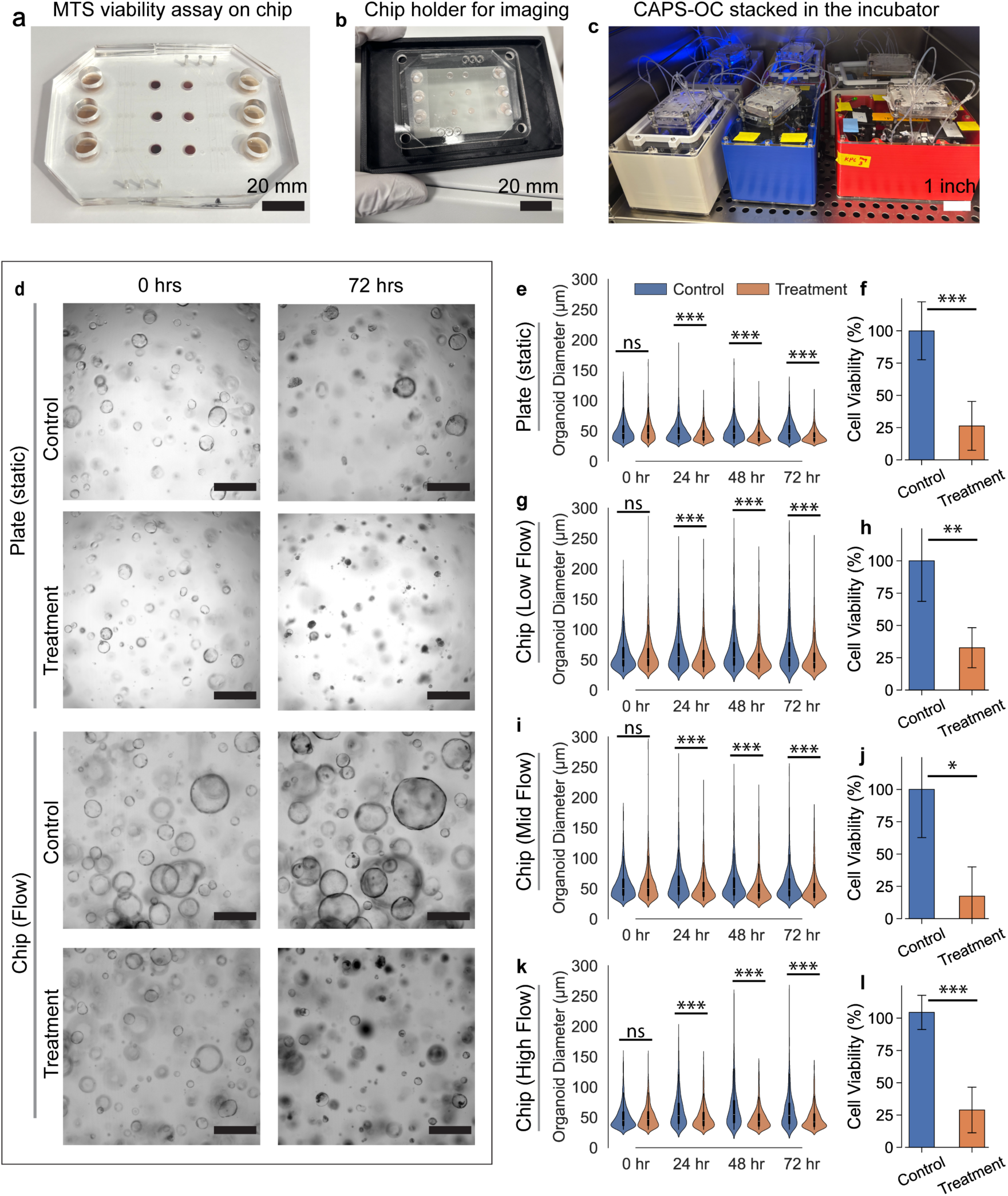
Drug response analysis of KPC organoids against multi-selective RAS inhibitor Daraxonrasib in the organoid-on-chip and conventional plate cultures. (a) MTS viability assay demonstrates that endpoint assessment of cell viability can be performed in a straightforward, user-friendly manner on-chip following drug treatment, providing quantitative readouts of metabolic activity. (b) Photograph of a custom 3D-printed microscope adapter that enables confocal imaging of organoids directly within the microfluidic chip, allowing longitudinal monitoring of growth and morphology without disturbing the culture. (c) Photograph showing multiple fully assembled organoid-on-chip units, illustrating the system’s scalability for medium-throughput drug dose–response studies. (d) Brightfield images comparing organoid morphology in response to a 72 hr incubation with RAS inhibitor-daraxonrasib, in both control and treatment groups, highlighting differences between plate and microfluidic chip cultures. Changes in organoid size and distributions were quantified to assess treatment effects. (e, g, i, k) Kernel density estimation plots of organoid diameter distributions under plate (e), low-flow (g), mid-flow (i), and high-flow (k) conditions, tracked every 24 hr over the 72 hr treatment period, capturing temporal dynamics of drug response across different flow regimes. (f, h, j, l) MTS-based cell viability measurements for control and treatment groups corresponding to plate (f), low-flow (h), mid-flow (j), and high-flow (l) conditions. Data demonstrates the combined effects of microfluidic flow and drug treatment on organoid viability. Statistical significance was determined using Welch’s t-test (n = 2; ns- not significant, *- P≤0.05, **- P≤0.01, ***- P≤0.001).

## 4. Conclusion & future work

In recent few years, bioengineering of 3D in-vitro systems has seen tremendous growth and these technologies are continuously increasing our understanding of tumor and its response to pharmacological drugs. Active and precision control of media is still a serious technical challenge despite several advancements in microfluidics. In this paper, by implementing a system level-based design approach, we demonstrate CAPS-OC enables culturing of PDAC organoids under a dynamic flow condition. Mainly, this system is entirely standalone and all-integrated without any peripheral components. We believe this system addresses several pain points of the existing flow-based microfluidic system and can be easily adopted by the bioengineering community. Using a clinically relevant model of KPC organoids, we show fluid flow is an influential microenvironment factor as it causes changes in size and morphology.

While the current version of the CAPS-OC prototype is effective, PDMS is not an ideal material for drug screening experiments. It is well established that small molecule drugs can leach into the PDMS substrate, requiring spiking of the dose to maintain consistent concentrations. While the current scope of work did not include this, we plan to develop a thermoplastic chip to circumvent this issue. Such chips can be made using layer-by-layer assembly of PMMA or PC bonded together via biocompatible pressure sensitive tapes. The chip design involving gas and fluid layer offered some challenges in terms of Matrigel seeding. In the current experimental protocol, we had to seed 12.5 µL gel in the 4 mm organoid wells in the fluidic layer followed by aligning of the gas layer under a microscope. This was feasible to demonstrate our proof-of-concept prototype but is certainly not user-friendly and is a rate limiting step. In the next version of the chip design, we would incorporate an “open organoid well” design with a removable gasketed insert. This way, users can simply seed the Matrigel into an open well and gasketed insert can seal it without blocking the flow path.

The current microfluidic chip is designed with a tangential flow over the encapsulating 3D matrix as opposed to a flow-through system. In this configuration, the pressure drop across the 3D Matrigel is negligible and as a result the interstitial flow and resulting shear stress observed by the organoids is expected to be negligible. The PDAC phenotypes we observed are thus likely because of diffusion of available nutrients cues, rather than hypoxic stress or global EMT. However, contribution of shear stress may not be totally ruled out without a scaling analysis and/or development of a flow-through chip.

Moreover, applications of our system are not limited to culturing organoids under flow since the chip design can be adapted to the application of interest. The current recirculatory loop design was used as the first proof of concept to show validity of the system. However, more biologically complex models can be built- for example, incorporation of vasculature along with organoids, fluidically connecting two organoids to study communication, so on and so forth.

## Acknowledgements

This work was supported in part by funding from the Cancer Early Detection Advanced Research Center at Oregon Health & Science University’s Knight Cancer Institute (Full 2023-1717). Alexander Davies acknowledges support from National Institutes of Health, Office of Research Infrastructure Programs, under award number K01OD031811. Luiz Bertassoni acknowledges support from NIH/NIDCR (R01DE035326) and NIH/NCI (R01CA310177). The research reported in this publication used computational infrastructure supported by the Office of Research Infrastructure Programs, Office of the Director, of the National Institutes of Health under Award Number S10OD034224. The content is solely the responsibility of the authors and does not necessarily represent the official views of the National Institutes of Health. Ellen Langer acknowledges support from a Research Scholar Grant, RSG-22-060-01-MM, grant https://doi.org/10.53354/pc.gr.153686, from the American Cancer Society. We would also like to thank Dr. Elie Traer, Keith Beadle, Mckenna Finley for useful discussions around design considerations and Madeline Kuhn for help with cell culture.

## References

1. Ingber, D. E. Challenges and opportunities for human Organ Chips in FDA assessments and pharma pipelines. Cell Stem Cell 33, 176–183 (2026).

2. Park, S. E., Georgescu, A. & Huh, D. Organoids-on-a-chip. Science (1979). 364, 960–965 (2019).

3. Liu, K. et al. From organoids to organoids-on-a-chip: Current applications and challenges in biomedical research. Chin. Med. J. (Engl). 138, 792 (2025).

4. Donoghue, L., Nguyen, K. T., Graham, C. & Sethu, P. Tissue chips and microphysiological systems for disease modeling and drug testing. Micromachines (Basel). 12, 1–35 (2021).

5. Mansouri, M., Lam, J. & Sung, K. E. Progress in developing microphysiological systems for biological product assessment. Lab Chip 24, 1293–1306 (2024).

6. Juste-Lanas, Y., Hervas-Raluy, S., García-Aznar, J. M. & González-Loyola, A. Fluid flow to mimic organ function in 3D in vitro models. APL Bioeng. 7, 031501 (2023).

7. Yang, J. et al. Dynamic culture system advances the applications of breast cancer organoids for precision medicine. Scientific Reports 2025 15:1 15, 8852- (2025).

8. Homan, K. A. et al. Flow-enhanced vascularization and maturation of kidney organoids in vitro. Nat. Methods 16, 255 (2019).

9. Shi, Z. D. & Tarbell, J. M. Fluid Flow Mechanotransduction in Vascular Smooth Muscle Cells and Fibroblasts. Ann. Biomed. Eng. 39, 1608 (2011).

10. Dessalles, C. A., Ramón-Lozano, C., Babataheri, A. & Barakat, A. I. Luminal flow actuation generates coupled shear and strain in a microvessel-on-chip. Biofabrication 14, 015003 (2021).

11. Quintard, C. et al. A microfluidic platform integrating functional vascularized organoids-on-chip. Nature Communications 2024 15:1 15, 1452- (2024).

12. Lee, K. K. et al. Human stomach-on-a-chip with luminal flow and peristaltic-like motility. Lab Chip 18, 3079 (2018).

13. Peristaltic Pumps: A Comprehensive Guide - Darwin Microfluidics. https://blog.darwin-microfluidics.com/peristaltic-pumps-a-comprehensive-guide/.

14. Ancelmo, H. C., Schuster da Silva, S., Kuchenbecker, H. F., Canniatti Brazaca, L. & Blanes, L. Microfluidic flow control strategies for organ-on-a-chip devices: a critical review. Discover Electronics 2026 3:1 3, 52- (2026).

15. Rajasekar, S. et al. IFlowPlate—A Customized 384-Well Plate for the Culture of Perfusable Vascularized Colon Organoids. Advanced Materials 32, 2002974 (2020).

16. Bonanini, F. et al. In vitro grafting of hepatic spheroids and organoids on a microfluidic vascular bed. Angiogenesis 2022 25:4 25, 455–470 (2022).

17. Chhabra, A. et al. A vascularized model of the human liver mimics regenerative responses. Proc. Natl. Acad. Sci. U. S. A. 119, e2115867119 (2022).

18. Lang, Q. et al. AC Electrothermal Circulatory Pumping Chip for Cell Culture. ACS Appl. Mater. Interfaces 7, 26792–26801 (2015).

19. Chang, J. Y. et al. A novel miniature dynamic microfluidic cell culture platform using electro-osmosis diode pumping. Biomicrofluidics 8, 044116 (2014).

20. Holman, J. B., Zhu, X. & Cheng, H. Piezoelectric micropump with integrated elastomeric check valves: design, performance characterization and primary application for 3D cell culture. Biomed. Microdevices 25, (2023).

21. Ma, T., Sun, S., Li, B. & Chu, J. Piezoelectric peristaltic micropump integrated on a microfluidic chip. Sens. Actuators A Phys. 292, 90–96 (2019).

22. Jeon, H. M., Kim, K., Choi, K. C. & Sung, G. Y. Side-effect test of sorafenib using 3-D skin equivalent based on microfluidic skin-on-a-chip. Journal of Industrial and Engineering Chemistry 82, 71–80 (2020).

23. Lee, D. W., Choi, N. & Sung, J. H. A microfluidic chip with gravity-induced unidirectional flow for perfusion cell culture. Biotechnol. Prog. 35, e2701 (2019).

24. Jeong, G. S. et al. Siphon-driven microfluidic passive pump with a yarn flow resistance controller. Lab Chip 14, 4213–4219 (2014).

25. Cha, K. J. & Kim, D. S. A portable pressure pump for microfluidic lab-on-a-chip systems using a porous polydimethylsiloxane (PDMS) sponge. Biomedical Microdevices 2011 13:5 13, 877–883 (2011).

26. Goral, V. N., Tran, E. & Yuena, P. K. A pump-free membrane-controlled perfusion microfluidic platform. Biomicrofluidics 9, 054103 (2015).

27. Pilarek, M., Neubauer, P. & Marx, U. Biological cardio-micro-pumps for microbioreactors and analytical micro-systems. Sens. Actuators B Chem. 156, 517–526 (2011).

28. Brower, K. et al. An open-source, programmable pneumatic setup for operation and automated control of single- and multi-layer microfluidic devices. HardwareX 3, 117–134 (2018).

29. de Graaf, M. N. S., Vivas, A., van der Meer, A. D., Mummery, C. L. & Orlova, V. V. Pressure-Driven Perfusion System to Control, Multiplex and Recirculate Cell Culture Medium for Organs-on-Chips. Micromachines (Basel). 13, 1359 (2022).

30. Boj, S. F. et al. Organoid models of human and mouse ductal pancreatic cancer. Cell 160, 324–338 (2015).

31. Lee, J. W., Komar, C. A., Bengsch, F., Graham, K. & Beatty, G. L. Genetically engineered mouse models of pancreatic cancer: The KPC model (LSL-KrasG12D/+;LSL-Trp53R172H/+;Pdx-1-Cre), its variants, and their application in immuno-oncology drug discovery. Curr. Protoc. Pharmacol. 2016, 14.39.1–14.39.20 (2016).

32. Hingorani, S. R. et al. Trp53R172H and KrasG12D cooperate to promote chromosomal instability and widely metastatic pancreatic ductal adenocarcinoma in mice. Cancer Cell 7, 469–483 (2005).

33. Low, R. R. J. et al. The Diverse Applications of Pancreatic Ductal Adenocarcinoma Organoids. Cancers 2021, Vol. 13, Page 4979 13, 4979 (2021).

34. Du, X. et al. Precision organoid segmentation technique (POST): accurate organoid segmentation in challenging bright-field images. Bio-Design and Manufacturing 2025 9:1 9, 80–93 (2025).

35. GitHub - duxuan11/Precision-Organoid-Segmentation-Technique-POST: segment any organoid · GitHub. https://github.com/duxuan11/Precision-Organoid-Segmentation-Technique-POST.

36. Bulle, A. & Lim, K. H. Beyond just a tight fortress: contribution of stroma to epithelial-mesenchymal transition in pancreatic cancer. Signal Transduct. Target. Ther. 5, 249 (2020).

37. Masugi, Y. The Desmoplastic Stroma of Pancreatic Cancer: Multilayered Levels of Heterogeneity, Clinical Significance, and Therapeutic Opportunities. Cancers (Basel). 14, 3293 (2022).

38. DuFort, C. C. et al. Interstitial Pressure in Pancreatic Ductal Adenocarcinoma Is Dominated by a Gel-Fluid Phase. Biophys. J. 110, 2106–2119 (2016).

39. Olive, K. P. et al. Inhibition of Hedgehog signaling enhances delivery of chemotherapy in a mouse model of pancreatic cancer. Science (1979). 324, 1457–1461 (2009).

40. Provenzano, P. P. et al. Enzymatic targeting of the stroma ablates physical barriers to treatment of pancreatic ductal adenocarcinoma. Cancer Cell 21, 418–429 (2012).

41. Fujita, H. et al. alpha-Smooth Muscle Actin Expressing Stroma Promotes an Aggressive Tumor Biology in Pancreatic Ductal Adenocarcinoma. Pancreas 39, 1254–1262 (2010).

42. Vonlaufen, A. et al. Pancreatic stellate cells: partners in crime with pancreatic cancer cells. Cancer Res. 68, 2085–2093 (2008).

43. Lahusen, A. et al. A pancreatic cancer organoid-in-matrix platform shows distinct sensitivities to T cell killing. Scientific Reports 2024 14:1 14, 9377- (2024).

44. Geyer, M. et al. A microfluidic-based PDAC organoid system reveals the impact of hypoxia in response to treatment. Cell Death Discov. 9, 20 (2023).

45. Kramer, B. et al. Interstitial Flow Recapitulates Gemcitabine Chemoresistance in A 3D Microfluidic Pancreatic Ductal Adenocarcinoma Model by Induction of Multidrug Resistance Proteins. Int. J. Mol. Sci. 20, (2019).

46. Toepke, M. W. & Beebe, D. J. PDMS absorption of small molecules and consequences in microfluidic applications. Lab Chip 6, 1484–1486 (2006).

47. Regehr, K. J. et al. Biological implications of polydimethylsiloxane-based microfluidic cell culture. Lab Chip 9, 2132–2139 (2009).

48. Auner, A. W., Tasneem, K. M., Markov, D. A., McCawley, L. J. & Hutson, M. S. Chemical-PDMS binding kinetics and implications for bioavailability in microfluidic devices. Lab Chip 19, 864–874 (2019).

49. Wang, J. D., Douville, N. J., Takayama, S. & ElSayed, M. Quantitative Analysis of Molecular Absorption into PDMS Microfluidic Channels. Ann. Biomed. Eng. 40, 1862–1873 (2012).

50. Torino, S., Corrado, B., Iodice, M. & Coppola, G. PDMS-Based Microfluidic Devices for Cell Culture. Inventions 2018, Vol. 3, Page 65 3, 65 (2018).

51. Hermann, N. G., Ficek, R. A., Markov, D. A., McCawley, L. J. & Hutson, M. S. Quantifying and modeling loss of estrogen and progesterone in PDMS-based devices. Microfluid. Nanofluidics 29, 78 (2025).

52. van Meer, B. J. et al. Small molecule absorption by PDMS in the context of drug response bioassays. Biochem. Biophys. Res. Commun. 482, 323–328 (2017).

53. Carter, S.-S. D. et al. PDMS leaching and its implications for on-chip studies focusing on bone regeneration applications. Organs-on-a-Chip 2, 100004 (2020).

54. Murthy, V. et al. Serial Imaging of Tumor and microEnvironment (SITE) platform for live-cell ex vivo modeling of primary and metastatic cancer dynamics. bioRxiv 2025.09.08.674915 (2025) doi:10.1101/2025.09.08.674915.

55. Sekiya, S., Kikuchi, T. & Shimizu, T. Perfusion culture maintained with an air-liquid interface to stimulate epithelial cell organization in renal organoids in vitro. BMC Biomed. Eng. 1, 15 (2019).

56. Cho, A. N. et al. Microfluidic device with brain extracellular matrix promotes structural and functional maturation of human brain organoids. Nature Communications 2021 12:1 12, 4730- (2021).

57. Martinelli, P. et al. GATA6 regulates EMT and tumour dissemination, and is a marker of response to adjuvant chemotherapy in pancreatic cancer. Gut 66, 1665–1676 (2017).

58. O’Kane, G. M. et al. GATA6 Expression Distinguishes Classical and Basal-like Subtypes in Advanced Pancreatic Cancer. Clin. Cancer Res. 26, 4901–4910 (2020).

59. Brunton, H., et al. *HNF4A* and *GATA6* Loss Reveals Therapeutically Actionable Subtypes in Pancreatic Cancer. Cell Rep. 31, 107625 (2020).

60. Duan, K. et al. The value of GATA6 immunohistochemistry and computer-assisted diagnosis to predict clinical outcome in advanced pancreatic cancer. Sci. Rep. 11, 14951 (2021).

61. Driehuis, E. et al. Pancreatic cancer organoids recapitulate disease and allow personalized drug screening. Proceedings of the National Academy of Sciences 116, 26580–26590 (2019).

