## Supplemental Figures for "A Compact, Standalone & Battery-Powered 3D Organoid-on-Chip System with Programmable Flow Control"

### Supplementary Information

#### Section 1: Printed circuit board

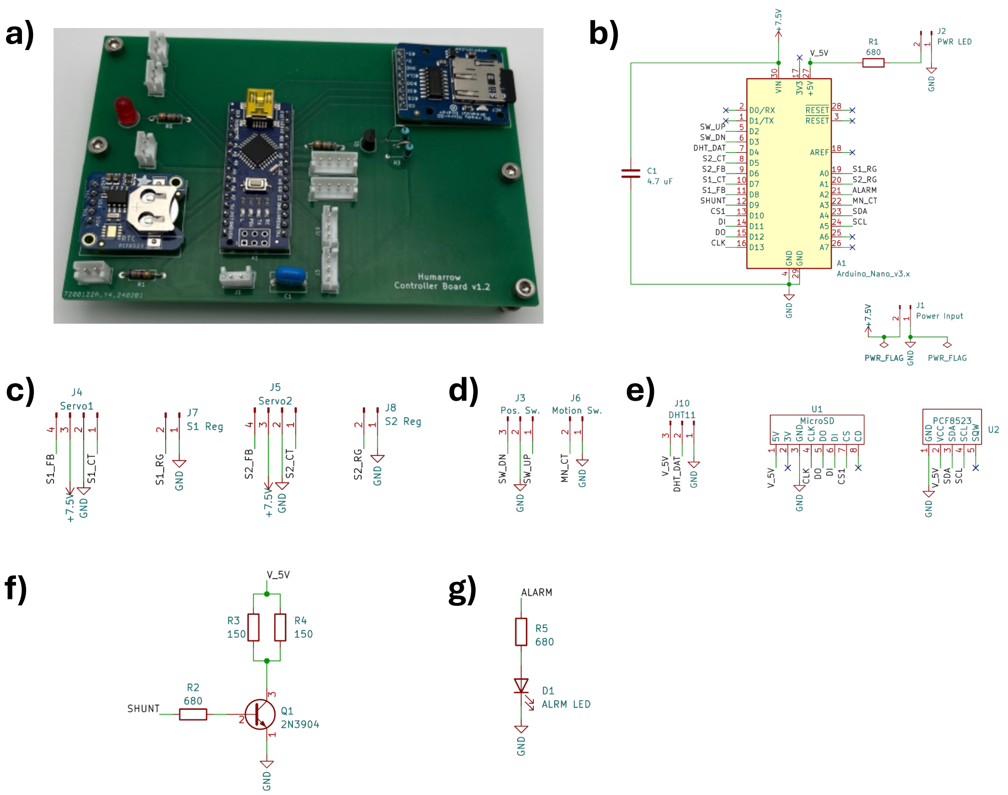


**Figure S1**: System driving electronics. a) Assembled printed circuit board. Electrical schematics for b) Arduino microcontroller circuit and power input, c) Servo power delivery and registration, d) user interface including positional switch and motion switch, e) data logging including a DHT11 temperature/humidity sensor, a MicroSD card reader/writer, and a real time clock, f) power shunt circuit, g) Alarm LED circuit

The driving electronics for the system are laid out on a custom printed circuit board (PCB) for ease of assembly (Fig. S1a). The control circuitry is based around an Arduino Nano microcontroller, which is responsible for digital and analog input and output, as well as state control (Fig. S1b). JST connectors are used to deliver power and communications from the Nano to the two servos, as well as to deliver limit switch signal back to the Nano for servo registration (Fig. S1c). The servo control signals (S1_CT and S2_CT) are connected to digital output pins on the Nano, which use pulse width modulation (PWM) to control servo movement. The servo feedback signals (S1_FB and S2_FB) are read by digital input pins on the Nano, where they are converted to a PWM modulation for determining servo angular location. The registration signals from the limit switches (S1_RG and S2_RG) are connected to digital input pins on the Nano that are pulled up through software using onboard pull-up resistors. When the limit switch is closed, the input pin is grounded allowing the Nano to detect that the servo has registered its position.

A 3-way position switch and a push-button motion switch are also connected to the PCB through JST connectors (Fig. S1d). The position switch allows the user to select the maximum clockwise position (when SW_DN is shorted to ground), the maximum counter-clockwise position (when SW_UP is shorted to ground), or at the midpoint between maximum clockwise and maximum counter-clockwise position (when neither SW_DN nor SW_UP are shorted to ground, i.e., the switch is in the middle position). The motion control button controls whether the movement is paused or continuous. If the button is depressed (i.e., MN_CT is shorted to ground), the movement is continuous and the 3-way position switch is ignored. If the button is not depressed, the movement is paused and the servos move to the position indicated by the 3-way position switch.

The system has the ability to log temperature and humidity data over time. This is enabled by the inclusion of a DHT11 temperature/humidity sensor, a MicroSD writer, and a real time clock on the PCB (Fig. S1e). The DHT11 is supplied 5 V and ground, and its data pin is connected to an analog input on the microcontroller (DHT_DAT). The MicroSD card reader also receives 5 V and ground from the microcontroller and has four additional pins connected to digital I/O pins on the Nano: clock (CLK), data out (DO), data in (DI), and chip select (CS1). The real time clock is also powered by the microcontroller, and additionally has its data (SDA) and clock (SCL) pins connected to digital I/O pins on the Nano.

A power shunt circuit is included on the PCB to prevent the power bank from restarting due to a low current draw when the servos are stationary (Fig. S1f). This shunt consists of a transistor switch controlled by a digital output pin on the microcontroller which, when open, causes 5 V to be applied across two 150 Ω resistors in series. When the switch is activated, a current of ~67 mA flows through the resistors and prevents the power bank from shutting off. The PCB also features an alarm LED powered by a digital output pin of the microcontroller which can be programmed to activate when a faulty state is encountered.

Section 2: Custom fabricated components

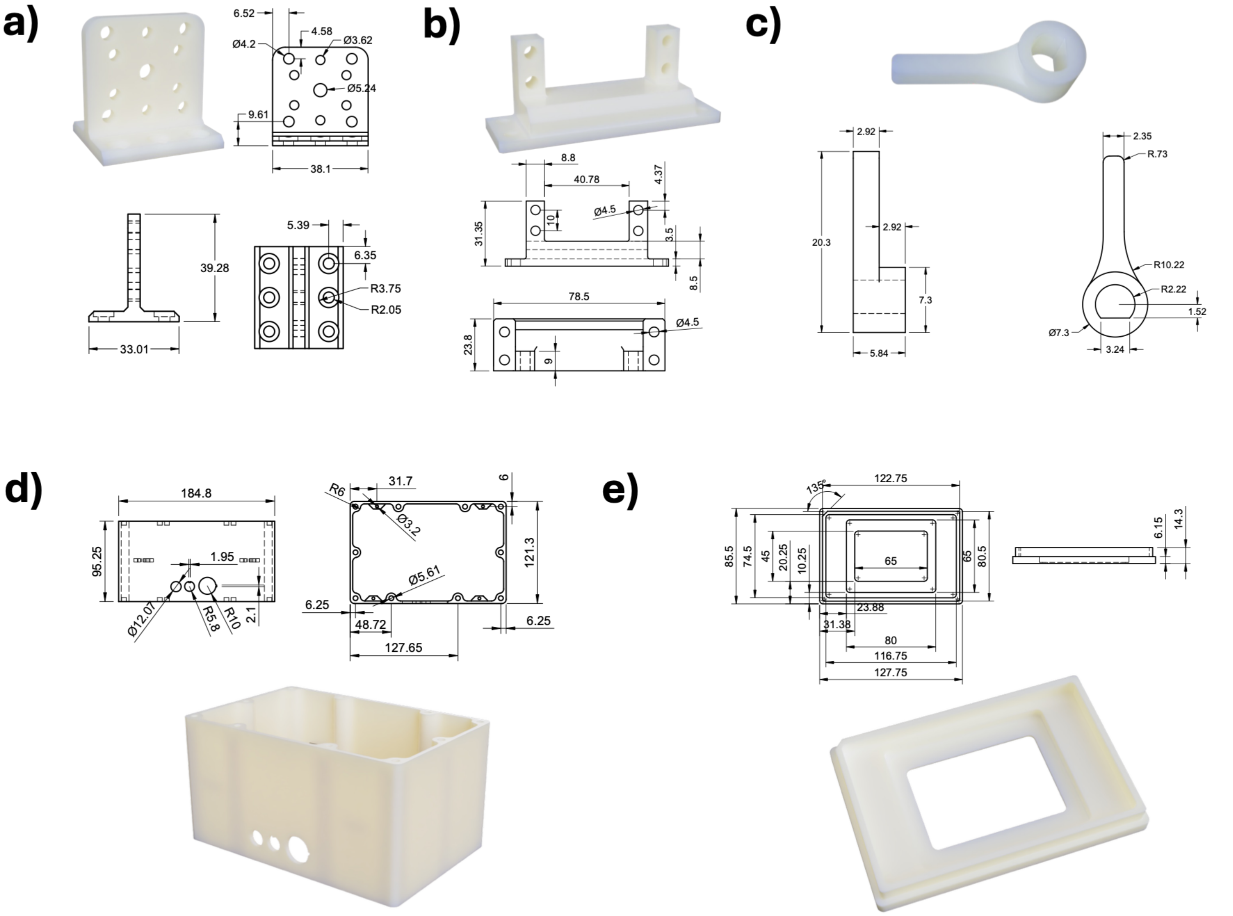


**Figure S2**: Drawings (units in mm) and renders of 3D-printed components: a) compact rotary actuator mount, b) servo motor mount, c) compact rotary actuator triggering arm, d) enclosure, e) microscope adapter

The system contains several custom-designed components which must be fabricated prior to assembly. The internal components and enclosure can be 3D-printed using the user's preferred material (suggested: PLA or ABS). The compact rotary actuator (CRA) mount, shown in Fig. S2a, holds the CRA in position to allow the driving servo to connect to it. The shaft of the CRA passes through the central 5.24 mm hole and is attached to the mount via four M3x10mm screws through the 4.2 mm holes on the corners of the mount. The mount is attached to the mid-plate of the system via six M4x14mm screws. The servomotor is also attached to the mid-plate via a 3D-printed mount, shown Fig. S2b. The servo attaches to the mount via four M4x20mm screws, and the mount is subsequently attached to the base with a second set of four M4x14mm screws.

A triggering arm must be added to the end of the CRA shaft to allow for interaction with the limit switches on the mid-plate for rotational registration. This arm, shown in Fig. S2c, is also 3D printed and simply slides onto the keyed end of the CRA shaft. The enclosure which holds all the internal components can also be 3D printed, and the design is shown in Fig. S2d. The dimensions of the box are 121.3 x 184.8 x 95.25 mm (WxLxH), and through-holes are included for the power switch, motion switch, and positional switch.

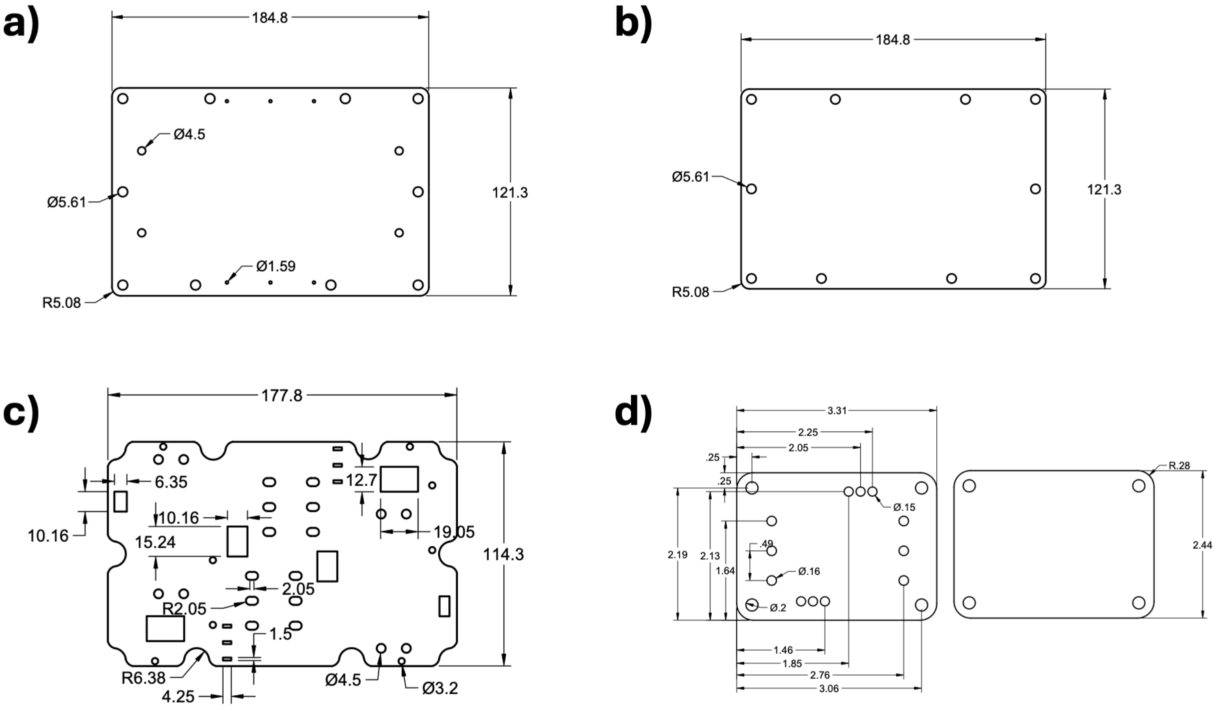


**Figure S3**: Drawings (units in mm) of laser-cut components, a) top plate, b) bottom plate, c) mid plate, d) chip clamping bracket. All plates are cut from 1/8” acrylic sheet

In addition to 3D-printed components, several components must be laser cut out of acrylic plastic. The top and bottom of the enclosure (Figs. S3a and S3b, respectively) help limit the entrance of moisture into the system. During experiments, these are further sealed using an elastomer membrane. The mid plate (Fig. S3c) sits in the middle of the enclosure and houses the 3D-mounted CRA and servo mounts, as well as the PCB. It includes through-holes for passing wires between the two sides of the enclosure.

#### Section 3: Assembly Instructions

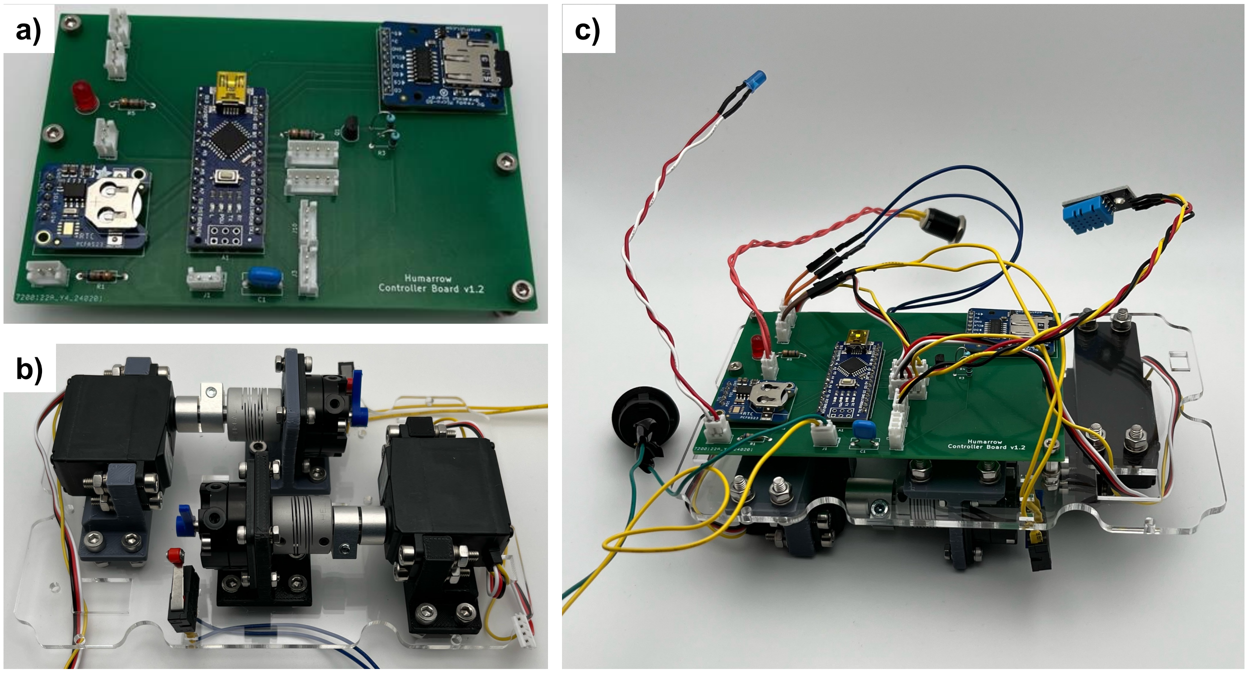


**Figure S4**: a) Assembled PCB, b) compact rotary actuators and servo motors attached to mid plate, c) PCB attached to underside of mid plate

##### *3.1 Preliminary Steps*

1. 3D-print components using 20% infill PLA or ABS using a 0.2 mm nozzle at 215 °C and a bed temperature of 60 °C
    a. Compact rotary actuator mount (x2)
    b. Servo motor mount (x2)
    c. Compact rotary actuator triggering arm (x2)
    d. Enclosure (x1)
2. Laser cut components from 1/8” clear acrylic (McMaster-Carr #8589K81) using a laser power of 45 W and a cutting speed of 15 mm/sec
    a. Top plate
    b. Bottom plate
    c. Mid plate
3. Obtain printed circuit board and solder components onto it following the circuit diagrams shown in Figure S1b-g, final assembly shown in Fig. S4a
    a. Solder headers onto the Arduino Nano (Arduino #ABX00028) and then solder it to the PCB at location A1
    b. Solder headers onto the MicroSD card module (Adafruit #254) and then solder it to the PCB at location U1
    c. Solder headers onto the real-time clock module (Adafruit #3295) and then solder it to the PCB at location U2
    d. Program the Arduino Nano with the PCF8523 example firmware in the Arduino IDE to set the time and date of the real-time clock module (File>Examples>RTClib>pcf8523)

e. Solder 680 Ω resistors (Digi-Key #CF14JT680RCT-ND) at locations R1, R2, and R5 on the PCB
f. Solder 150 Ω resistors (Digi-Key #RNF14FTD150RCT-ND) at locations R3 and R4 on the PCB
g. Solder a 2N3904 NPN transistor (Digi-Key #4491-2N3904-ND) at location Q1 on the PCB
h. Solder a 4.7 µF ceramic capacitor (Digi-Key #445-181580-1-ND) at location C1 on the PCB
i. Solder a red LED (Digi-Key #365-1182-ND) at location D1 on the PCB
j. Solder 2-pin JST receptacles (Digi-Key #455-B2B-EH-A-ND) at locations J1, J2, J6, J7, and J8 on the PCB
k. Solder 3-pin JST receptacles (Digi-Key #455-B3B-EH-A-ND) at locations J3 and J10 on the PCB
l. Solder 4-pin JST receptacles (Digi-Key #455-B4B-EH-A-ND) at locations J4 and J5 on the PCB

*3.2 Assembly of mechanical components*

1. Attach the 3D-printed servo motor mounts to the mid plate using four M4x14mm screws (McMaster-Carr #91292A038) with washers (McMaster-Carr #98689A113), lock washers (McMaster-Carr #94241A520), and nuts (McMaster-Carr #91828A231), as in Fig. S4b
2. Attach the 3D-printed CRA mount to the mid plate using six M4x12mm screws (McMaster-Carr #91292A117) with washers (McMaster-Carr #98689A113), lock washers (McMaster-Carr #94241A520), and nuts (McMaster-Carr #91828A231)
3. Using wire cutters, clip the female headers off the servo motor wires
    a. Strip ~1 mm from each wire jacket and tin the exposed braided wire with a small amount of solder
    b. Place a JST crimp (Digi-Key #455-1042-1-ND) onto the end of each wire
    c. Insert the wires into a 4-pin JST header (Digi-Key #455-1002-ND) per the following scheme:
        i. Position 1: white (control)
        ii. Position 2: black (ground)
        iii. Position 3: red (power)
        iv. Position 4: yellow (feedback)
4. Using a vise, press a 25-tooth spline to 1/4" round bore shaft coupler (ServoCity #4001-0025-0250) onto the shaft of the servo motor
5. Insert a 1/4" diameter, 3/4" long metal dowel rod (McMaster-Carr #98381A537) into the shaft coupler as far as it will go and tighten the collar to secure it in place
6. Attach a flexible shaft coupler (McMaster-Carr #6208K431) to the other end of the metal dowel leaving ~1 mm of space and tighten the collar to secure it in place
7. Attach the servo (Parallax #900-00360) to the 3D-printed servo mount using four M4x20mm screws (McMaster-Carr #91292A121, with washers (McMaster-Carr #98689A113), lock washers (McMaster-Carr #94241A520), and nuts (McMaster-Carr #91828A231)
8. Secure a flange bracket (McMaster-Carr #6508K42) to the CRA

9. Attach the CRA to the 3D-printed CRA mount using four M3x10mm screws (McMaster-Carr #91292A113), lock washers (McMaster-Carr #92148A150), and nuts (McMaster-Carr #91828A211)
10. Place a 3D-printed CRA triggering arm onto the keyed CRA shaft
11. Attach the other end of the CRA shaft to the flexible shaft coupler, connecting it to the servo motor
12. Solder wires to the two “normally open” leads of the limit switches (Amazon #B07X142VGC)
13. Thread the wires of the limit switches through the small slots in the mid plate with the arms positioned closer to the adjacent CRA
14. Attach JST crimps (Digi-Key #455-1042-1-ND) to the ends of each of the limit switch wires
15. Attach the assembled PCB to the underside of the mid plate using four M3x18mm screws (McMaster-Carr #91292A029), 6mm standoffs (McMaster-Carr #94669A101), and M3 nuts (McMaster-Carr #91828A211), as in Fig. S4c
16. Attach the JST connectors of the servo motors and the limit switches to their corresponding locations on the PCB

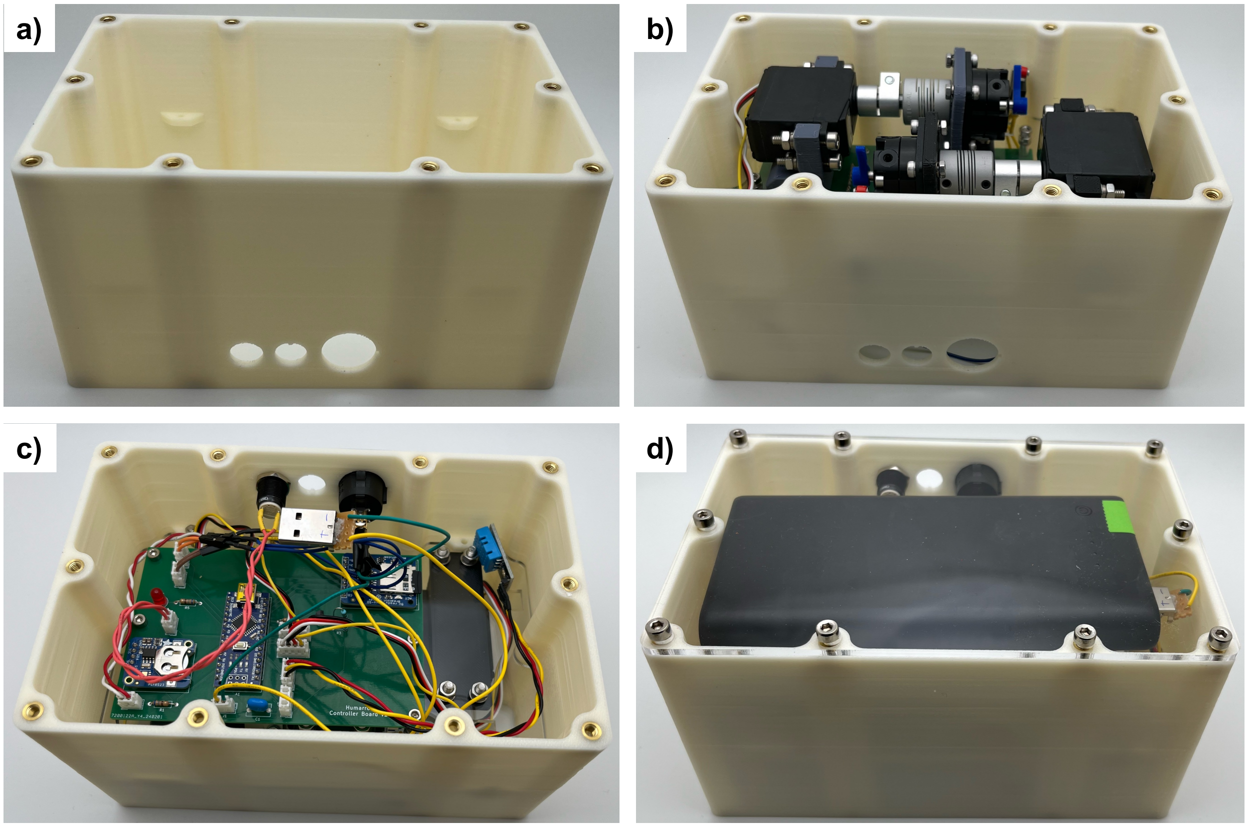


**Figure S5**: a) Enclosure with M4 brass threaded inserts, b) enclosure with mid plate added, c) underside of mid plate, showing PCB and user interface attached via JST connectors, d) battery added and bottom plate attached

*3.3 Assembly of the enclosure*

1. Using a soldering iron, heat set M4x4.7mm brass threaded inserts (McMaster-Carr #94180A351) into the 5 mm diameter holes in the top (x10) and bottom (x10) of the enclosure (Fig. S5a)
2. Place the mid plate in the enclosure and secure it with four M3x10mm screws (McMaster-Carr #91292A113) and M3 nuts (McMaster-Carr #91828A211), as in Fig. S5b
3. Solder wires to a 3-position switch (Amazon #B01J31IBH0), add JST crimp tins to the other side and insert them into a 3-position JST receptacle (Digi-Key #455-1001-ND), and insert the assembly into the 12 mm keyed hole on the front of the enclosure
4. Solder wires to a push-button switch (Amazon #B0C2GBN5MX), add JST crimp tins to the other side and insert them into a 2-position JST receptacle (Digi-Key #455-1000-ND), and insert the assembly into the 12 mm circular hole on the front of the enclosure
5. Assemble the power switch (Amazon #B07S1MV462)
    a. Solder one end of a wire to a terminal of the power switch and the other end of the wire to the positive terminal of a male USB receptacle (TinkerSphere #TS2532)
    b. Solder a wire to the ground terminal of the male USB receptacle and place the other end in one position of a 2-position JST connector (Digi-Key #455-1000-ND)
    c. Solder a wire to the other terminal of the power switch and place it in the other position of the 2-position JST connector
    d. Place the power switch with soldered USB receptacle in the 20 mm keyed hole in the front of the enclosure
6. Attach the JST connectors for the 3-position switch, push-button switch, and power switch to their corresponding receptacles on the PCB (Fig. S5c)
7. Connect a 26,800 mAh power bank (Amazon #B07ZX22KJS) to the male USB receptacle and place the power bank in the enclosure below the mid plate
8. Attach the bottom plate to the enclosure using 10 M4x6mm screws (McMaster-Carr #91292A107), as in Fig. S5d
9. Attach the chip holder to the top plate using 4 M3x16mm screws (McMaster-Carr #91292A115)
10. Attach the top plate with chip holder to the enclosure using 10 M4x6mm screws (McMaster-Carr #91292A107)

#### Section 4: Validation

**Supplementary Table 1**: PDMS membrane thickness measurements

| | Thickness (μm) | | | |
|---|---|---|---|---|
| **n** | **Strip 1** | **Strip 2** | **Strip 3** | **Strip 4** |
| **1** | 105.9 | 98.2 | 127.3 | 112 |
| **2** | 110.9 | 116.3 | 135.3 | 121.6 |
| **3** | 110.6 | 103.5 | 144.7 | 129.9 |
| **Avg. +/- S.D.** | 109.1 +/- 2.8 | 106.0 +/- 9.3 | 135.8 +/- 8.7 | 121.2 +/- 9.0 |

**Supplementary Table 2:** Time between strokes for various duty cycles (D.C.)

| | Delay time (ms) | | |
|---|---|---|---|
| **n** | **1% D.C.** | **10% D.C.** | **95% D.C.** |
| **1** | 294723 | 25263 | 139 |
| **2** | 290466 | 25056 | 142 |
| **3** | 290466 | 25254 | 157 |

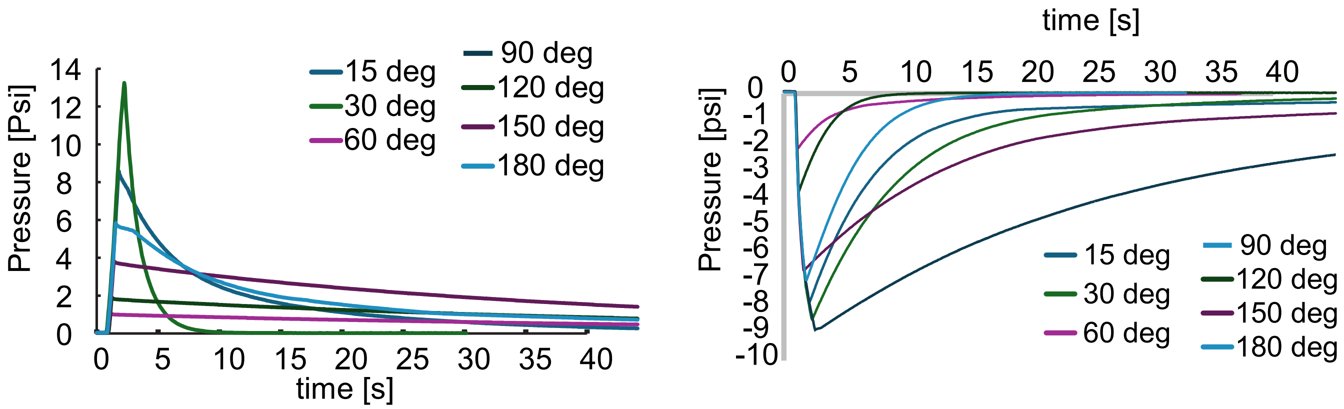


**Figure S6**: Example plots showing dissipation for different servo turn angles for clockwise (left) and for anticlockwise (right) angles.

#### Section 6: Biological Experiments

**Supplementary Table 3:** Mouse organoid splitting medium

| **component** | **Catalog number** | **Stock conc.** | **Final conc.** | **volume** |
|---|---|---|---|---|
| Advanced DMEM/F12 | 12634010 | | | 485 mL |
| HEPES pH 7.2-7.5 | 15630106 | 1 M | 10 nM | 5 mL |
| Penicillin-streptomycin | 15140122 | 100X | 1X | 5 mL |
| GlutaMAX | 35050061 | 100X | 1X | 5 mL |

**Supplementary Table 4:** Mouse organoid culture medium

| component | Catalog number | Stock conc. | Final conc. | Volume for 20 mL total |
|---|---|---|---|---|
| L-WRN conditioned medium | | | | 10 mL |
| RSPO-1 conditioned medium | | | | 2 mL |
| Advanced DMEM/F12 | 12634010 | | | 7.5 mL |
| GlutaMAX | 35050061 | 100X | 1X | 80 µL |
| B27 | 17504044 | 50X | 1X | 400 µL |
| N-acetylcysteine | A9165-25G | 500 mM | 1 mM | 40 µL |
| Nicotinamide | N0636-100G | 1M | 10 mM (1.22 mg/mL) | 200 µL |
| A83-01 | 9001799 | 0.5 mM (0.21 mg/mL) | 0.5 µM (0.21 ug/mL) | 20 µL |
| mEGF | 315-09 | 50 ug/mL | 0.05 µg/mL | 20 µL |
| SB202190 | 10010399 | 10 mM | 10 µM | 20 µL |
| PGE2 | 14010 | 0.5 mM | 500 nM | 20 µL |

**Supplementary Table 5**: **Statistical analysis of KPC organoid volumes within conditions across timepoints.**
Pairwise comparisons of organoid volumes were performed using Kruskal-Wallis test with Dunn's post-hoc analysis and Bonferroni correction. Significance is indicated as *-P<0.05, **-P<0.01, ***-P<0.001, ns=Not significant.

| | Condition | Significance |
|---|---|---|
| Static | Day 1 vs Day 3 | *** |
| | Day 1 vs Day 5 | *** |
| | Day 3 vs Day 5 | ns |
| Low Flow | Day 1 vs Day 3 | *** |
| | Day 1 vs Day 5 | *** |
| | Day 3 vs Day 5 | ns |
| Mid Flow | Day 1 vs Day 3 | *** |
| | Day 1 vs Day 5 | *** |
| | Day 3 vs Day 5 | * |
| High Flow | Day 1 vs Day 3 | *** |
| | Day 1 vs Day 5 | *** |
| | Day 3 vs Day 5 | ns |

**Supplementary Table 6**: **Statistical analysis of KPC organoid volumes across conditions and timepoints.**
Pairwise comparisons of organoid volumes were performed using Kruskal-Wallis test with Dunn's post-hoc analysis and Bonferroni correction. Significance is indicated as *-P<0.05, **-P<0.01, ***-P<0.001, ns=Not significant.

| | Condition | Significance |
|---|---|---|
| Day 1 | Static vs Low Flow | * |
| | Static vs Mid Flow | ns |
| | Static vs High Flow | ns |
| | Low Flow vs Mid Flow | ns |
| | Low Flow vs High Flow | ns |
| | Mid Flow vs High Flow | ns |
| Day 3 | Static vs Low Flow | *** |
| | Static vs Mid Flow | *** |
| | Static vs High Flow | * |
| | Low Flow vs Mid Flow | ns |
| | Low Flow vs High Flow | ** |
| | Mid Flow vs High Flow | * |
| Day 5 | Static vs Low Flow | ** |
| | Static vs Mid Flow | *** |
| | Static vs High Flow | ns |
| | Low Flow vs Mid Flow | ns |
| | Low Flow vs High Flow | * |
| | Mid Flow vs High Flow | *** |

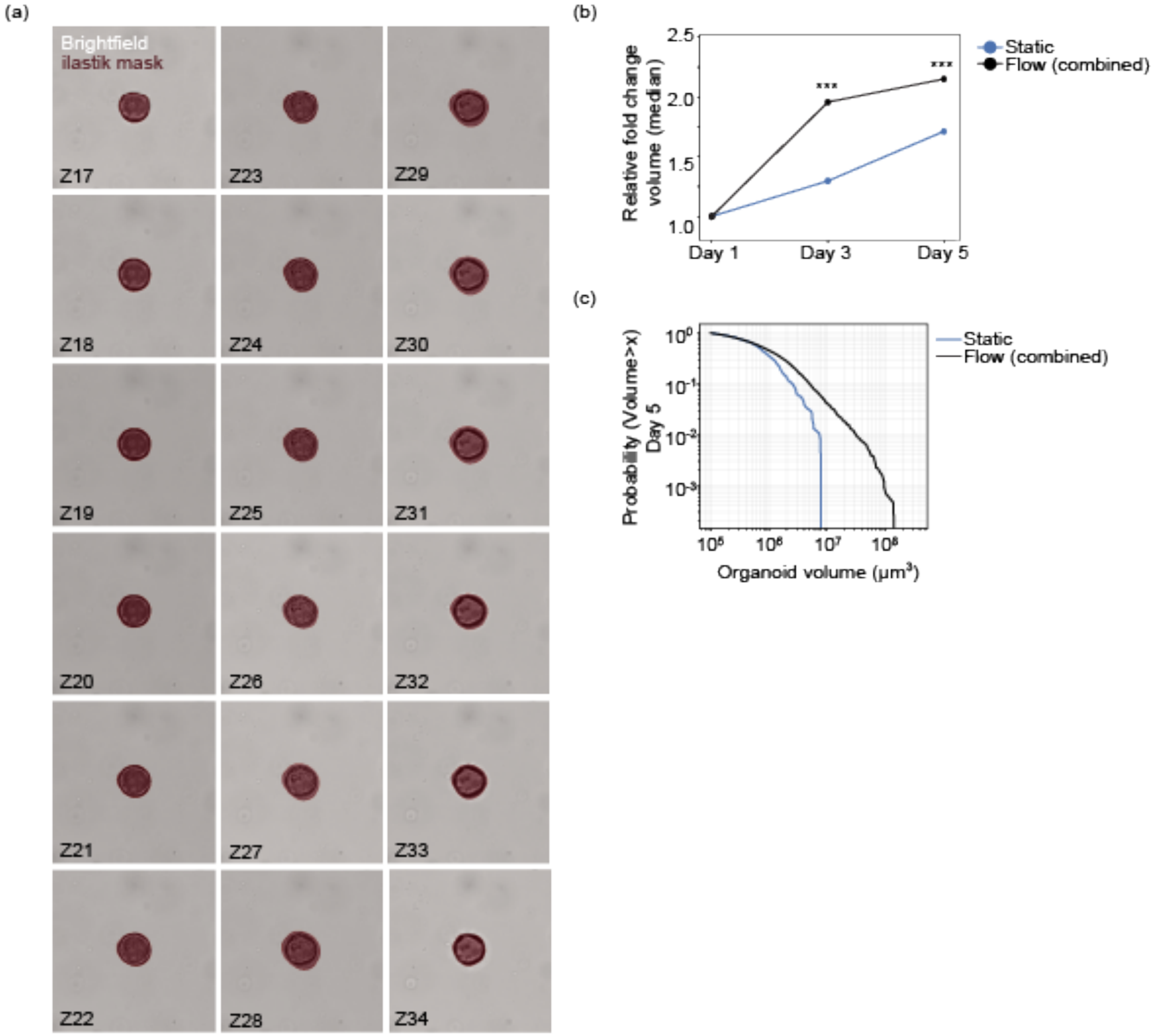


**Figure S7**: (a) Subset of individual Z-stack slices showing segmentation of ilastik masks (red) overlayed on brightfield (grey) KPC organoids. Segmentation was performed using machine learning approach (ilastik, N=50 training images). (b) Line plots showing relative fold change in organoid volume from day 1 in static (blue) and flow (combined, black) conditions. Significance was determined by means of a Welch's t-test (***- P≤0.0001) on independent experiments. (c) Complementary cumulative distribution function (CCDF) of KPC organoid volumes under static (blue), and flow (combined, black) conditions at day 5. The y-axis represents the probability that a randomly selected organoid exceeds a given volume threshold (x-axis, log scale).

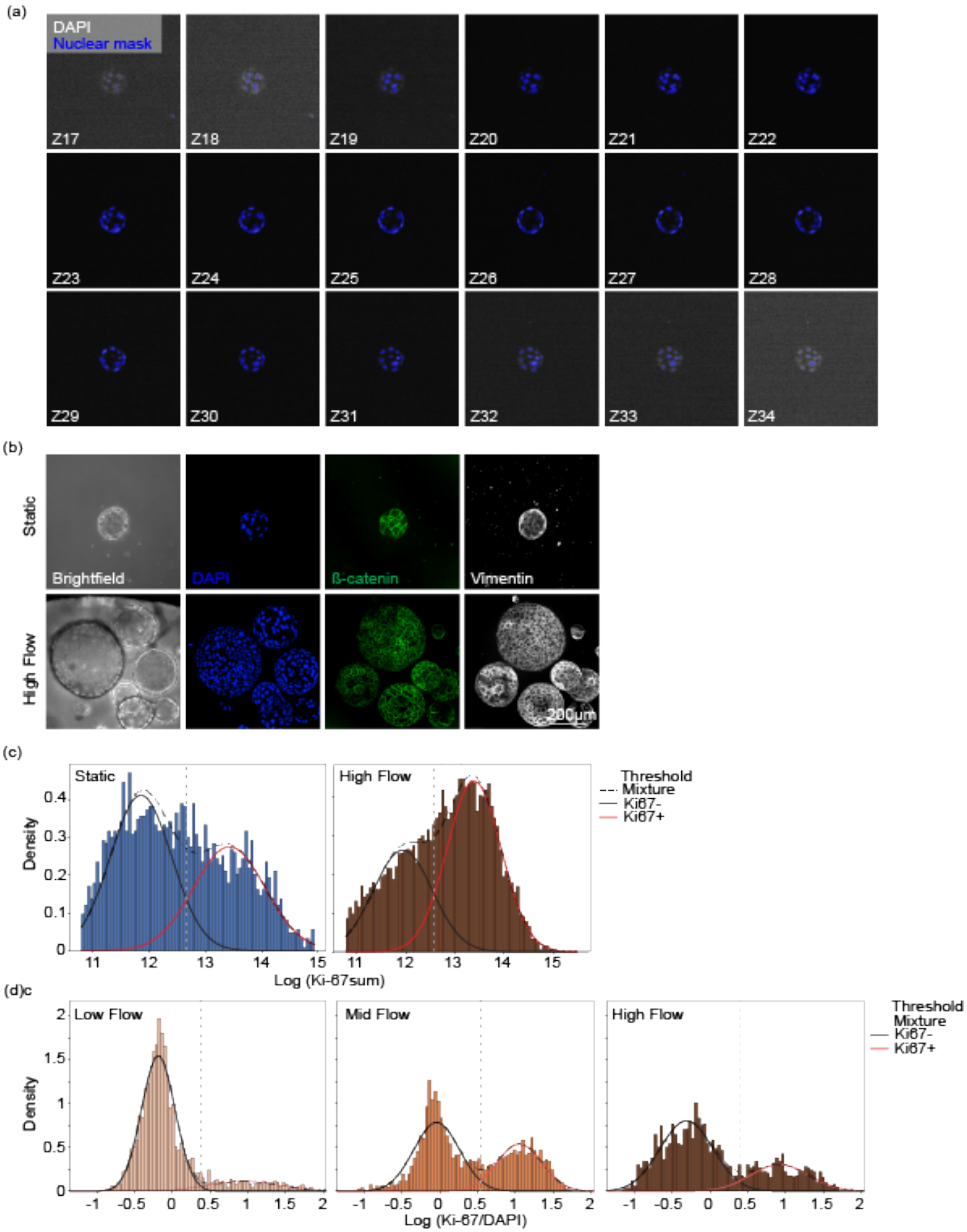


**Figure S8**: (a) Subset of individual Z-stack slices showing segmentation of nuclei (blue) overlayed on DAPI (grey) of KPC organoids. Segmentation was performed using machine learning approach (Cellpose, N=500 training images). (b) Representative maximum intensity projection images of Brightfield (grey), DAPI (blue), β-Catenin (green), and vimentin (white) in static and high flow conditions imaged at day 5.

Scale bar=200μm. (c) Two-component Gaussian mixed model fitted to the log-transformed total Ki67 sum intensity per nucleus for static (left) and high flow (right) conditions. (d) Two-component Gaussian mixed model fitted to the log-transformed Ki67/DAPI ratio for low flow (left), mid flow (middle), and high flow (right) conditions. Blue and red curves represent the Ki67− and Ki67+ Gaussian components respectively, the dashed black line shows the mixture distribution, and the grey dotted vertical line indicates the GMM-derived classification threshold.
